# Testing methodological and population-specific influences on the detection of Zipf’s law of brevity in chimpanzee gestures

**DOI:** 10.1101/2025.11.21.689682

**Authors:** A Safryghin, G Badihi, R Ferrer-i-Cancho, C Grund, M Hayashi, A Mielke, J Mine, ED Rodrigues, A Soldati, K Zuberbühler, C Zulberti, C Hobaiter

**Affiliations:** School of Psychology and Neuroscience, University of St Andrews, St Andrews,UK; School of Biological and Behavioural Sciences, Department of Psychology, Queen Mary University of London; Quantitative, Mathematical and Computational Linguistics Research Group, Departament de Ciències de la Computació, Universitat Politècnica de Catalunya, 08034 Barcelona, Catalonia, Spain; Chubu Gakuin University, Kakamigahara, Japan; Japan Monkey Centre, Inuyama, Japan; Department of Animal and Human Ethology, University of Rennes, Rennes, France; William James Center for Research, ISPA – Instituto Universitário, Lisbon, Portugal; Católica Research Centre for Psychological-Family and Social Wellbeing, Universidade Católica Portuguesa, Lisboa, Portugal; Department of Paleontology, University of Zurich, Zurich, Switzerland; Institute of Biology, University of Neuchâtel, Neuchâtel 2000, Switzerland; Institute of Biology, University of Leipzig, Leipzig, Germany

**Keywords:** compression, communication, linguistic law, ape, gesture

## Abstract

Zipf’s law of brevity is a widespread manifestation of information compression found across human languages and other communication systems. Chimpanzee gesture represents a rare absence of its expression in short-range communication. But, whether this absence reflects a feature of ape gesture production, or results from methodological obstacles or population differences is unclear. Selecting the appropriate unit of analysis is crucial for detecting linguistic patterns in any system. We assess gestural repertoires of three chimpanzee communities for Zipf’s law, while discriminating and testing different levels of unit durations (segmentation) and how finely gestural units are split (granularity). We report the first repertoire-wide detection of Zipf’s law in ape gesture in one chimpanzee community at a specific level of segmentation and granularity. We suggest that both methodological and socio-ecological factors can shape the detection and expression of Zipf’s law, emphasizing the importance of species-relevant units and metrics for meaningful cross-species comparisons.

## Introduction

‘Language laws’ are statistical regularities found across human languages. Zipf’s law of brevity (hereafter *Zipf’s law*) represents the tendency for more frequently used words to be shorter in length [1–2] and is generalised as the tendency for more frequently used elements of many kinds (e.g., syllables, words, calls) to be shorter or smaller [3]. Found in spoken, written, and signed languages [3–5], Zipf’s law is argued to represent a manifestation of the information theoretic principle of compression [6–7]. Based on the principle of least effort [2], Zipf’s law promotes coding efficiency by reducing the energy needed to produce a communicative code while retaining the information that needs to be transmitted [7–8].

First described in human languages, Zipf’s law has been documented across communicative systems including some bird songs (e.g., penguin display songs; [9], Western tanager songs [10]), primate and other mammalian vocalisations (e.g., Formosan macaques, [11]; Rock hyraxes, [12]; bats [13], whale songs [14–16], primate calls – for a review see [17]) and has been proposed as a widespread biological—rather than strictly linguistic—principle [6]. In human languages, including sign languages, Zipf’s law holds across levels of linguistic segmentation (e.g., when measured by duration, number of characters) and units (e.g., words, phonemes, signs, fingerspelling) [1, 5, 18–19]. In contrast, while phylogenetically widespread, nonhuman animal (hereafter ‘animal’) communicative systems show more variation in the units in which Zipf’s law is detected.

Language likely emerged in face-to-face interactions [20], and other species similarly show Zipf’s law as typically present in short range or close calls, but absent in some long-range vocalisations (bat species, [13, 21]; Rock hyrax, [12]; frogs, [22]; bird species [10], although see [16, 23]). In other species, its expression appears to be sensitive to different measures of energetic ‘investment’—for example, it is found in unit duration in some systems (Indri songs, [24]) and in call amplitude, but not duration, in others (Rock hyrax vocalisations, [12]). Ape gestural communication offers an ideal model system for comparison with language—well-described, it shares key properties with linguistic communication, such as intentionality, flexibility, and the combination of very large sets of signals into sequences [25–30]. However, recent work to investigate Zipf’s law in ape gesture revealed that it represented a rare exception in short-range communication: no evidence of Zipf’s law was detected in chimpanzees’ gesturing [31–32]. Given the proposed universality of the law [6] and its widespread presence across communicative systems - albeit represented through different units of energetic investment - any absence of Zipf’s law remains a puzzling exception to the rule that Zipf’s law of brevity seems to be a default across communicative systems. As a result, the apparent absence of Zipf’s law in ape gesture warrants further investigation: does it represent a true absence of the pattern, or have we, to date, been looking in the wrong place?

One explanation for the repertoire-level absence of Zipf’s law in previous studies may be found in the contexts of signal production. The first study investigated chimpanzee gesturing during play [31]. Gestural communication in play typically contains the majority of the very large repertoire of ape gesture actions, and the frequent use of sequences by a wide range of individuals [32–35], offering an apparently ideal dataset. However, play also reflects a context in which there may be relatively limited pressure towards efficient communication, serving as a practice ground for immature individuals [29] and an important means to foster social bonds throughout life [35]. A subsequent study investigated Zipf’s law in the gestural sequences produced during sexual solicitations [32], which also contain a wide range of gesture types and extensive use of sequences [31, 34]. Here, perhaps, the pressure towards successful communication may counteract any pressure towards brevity. In courtship, a small additional investment in signalling may ensure that signals are not missed or misunderstood, increasing probabilities of mating success. Moreover, individuals may invest in energetically demanding displays to advertise fitness. Hence, courtship signalling may be resistant to selection on energetically-efficient communication [32]. Finally, both studies were conducted on the same community of Eastern chimpanzees (*Pan troglodytes schweinfurthii*)—whose group-specific socio-ecology may shape communication in ways that do not generalise to chimpanzees as a species [38–39]. Thus, while the study of Zipf’s law in other species has been similarly limited to specific social contexts and populations [9, 12, 23], it remains possible that in ape gesture we have been measuring units in contexts where efficiency may have not exerted a strong selective pressure. The addition of further communities, populations, and contexts provides one means to address whether the apparent absence of Zipf’s law in ape gesture is generalisable at a species-level.

In addition, the detection of linguistic laws can be sensitive to the ways in which researchers quantify communicative units [40]. For instance, the emergence of another linguistic pattern—Zipf’s law for word frequencies [2]—depends on how words are categorised. This law is present when defining words as strings separated by spaces or punctuation marks, or as their ‘lemma’ (e.g., lemma of ‘swims’ and ‘swimming’ is ‘swim’ [41]) but absent when using a broader classification of words as their part-of-speech category (noun, verb, adjective, adverb, etc.) [42], or when focusing on individual letters [43]. Similarly, Andres et al. [43] initially failed to find Menzerath-Altmann’s law (describing the relationship between the number of parts of a sign and the size of its parts), in Czech sign language; but later detected it by re-clustering signs into pseudosyllables (correlates of spoken syllables).

In previous work on Zipf’s law in chimpanzee gestures, the gesture units analysed may not have been the ones in which Zipf’s law is typically expressed. Unlike vocal systems, in which there are clear limitations on call duration imposed by breathing, after a gestural action is produced it can be held in place for a substantial amount of time. Imagine an outstretched palm in a ‘Reach’ gesture, held in place until the requested item is handed over—a similar vocal request would not be held in place for the same duration while waiting for a response. The initial part of the gesture action—like the vocal unit—is more consistently produced, but the duration of the hold – largely absent in vocalisations – may be highly variable, depending on the specific set of communicative circumstances. In gestural systems, selective pressure towards efficient production may act differently on different parts of a gestural unit.

In human languages, units can be defined based on different levels of *granularity* (e.g., phrases, words, morphemes, phonemes). What is the ‘core’ of a gesture unit? Contrary to the study of human languages, an understanding of the parts making a call or gesture in other species is still in its infancy [45–46]. Early animal communication work took a top-down approach: human observers segmented units according to both their own perception and, often, the data available. But, this approach can lead to a structurally-anthropocentric bias that would limit our assessment of relevant unit of measure from a particular species’ perspective. With larger datasets now available, recent work has taken more of a ‘bottom-up’ approach, describing units at multiple levels of segmentation within a dataset [47–48]. In gesture, each unit can be segmented into discrete phases, which typically include periods of preparation and action stroke, and optional periods of hold, repetition, and/or recovery of the gesture action [40]. The preparation and gesture stroke can be grouped into the Minimum Action Unit (MAU, i.e. the shortest part of the gesture necessary to discriminate one type of gesture action from another; cf. [48]), and distinguished from the additional, optional production of a hold or repetition phase (Figure 1). Together with the MAU this extended expression of the gesture is termed the Performed Action Unit (PAU, Figure 1).

**Figure 1.**
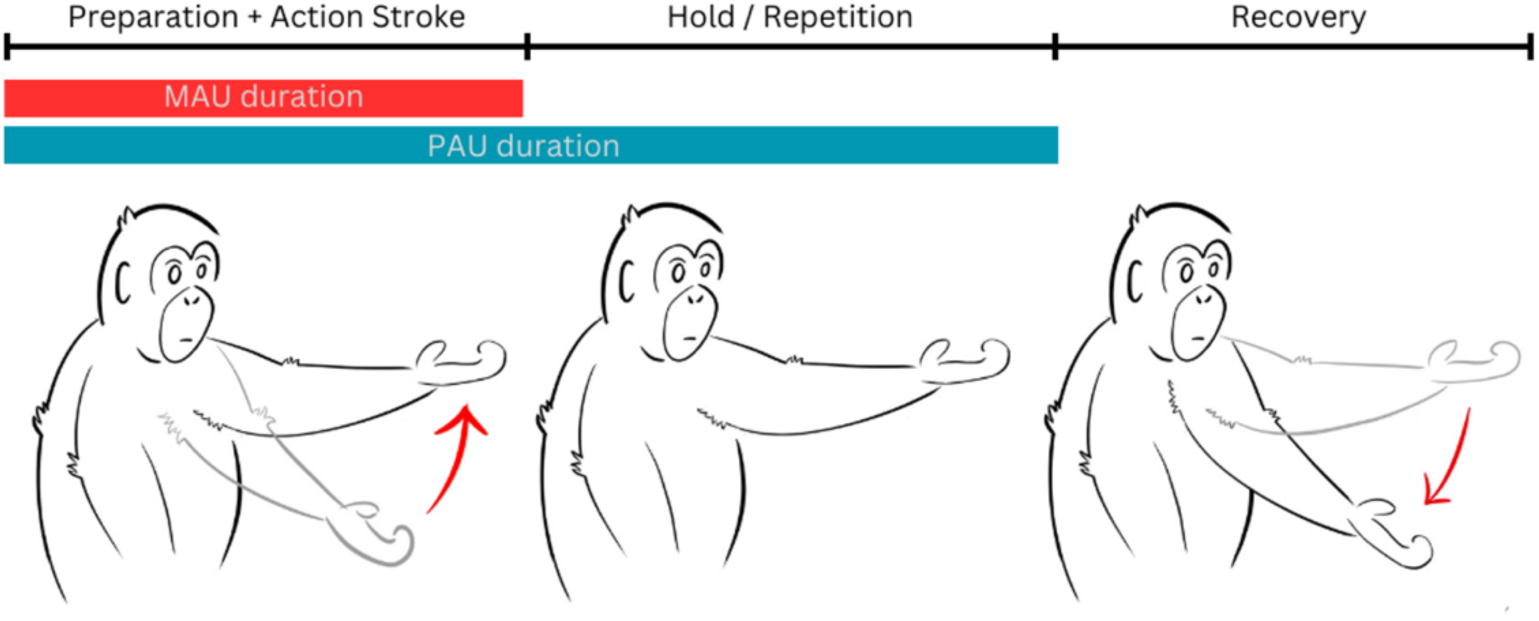
Gesture phases and gesture part durations for the Gesture Action *Reach*. The Minimum Action Unit (MAU) duration is represented by the red line, while the Performed Action Unit (PAU) duration is represented by the blue line.

Given that Zipf’s law operates at the level of signal ‘types’ in a repertoire (e.g., words in a lexicon), with more frequently-used signals being shorter and more infrequently-used signals longer, the duration of the MAU—present in every instance in which a gesture unit of that type is produced—may be more likely to be subject to generalised selection for compression. In contrast, the production and duration of an additional hold or repetition phase, is more likely to be sensitive to the immediate context of each communication generating substantial variation in total gesture duration [48–49]. As in the *Reach* example earlier, socio-ecological circumstances, pragmatic context, recipient responsiveness, or gesture position in a sequence all impact the likelihood with which a signaller ‘holds’ or ‘repeats’ the movement of a gesture in any particular instance [49]. Previous studies of Zipf’s law in chimpanzee gesture included these optional aspects of gesture production (i.e., the PAU and the recovery phases), whichmay have added substantial noise, hindering the detection of system-wide patterns of compression.

Moreover, as is the case in human languages, where units can be defined based on different levels of *granularity*, (as above, the lemma: swim, vs the forms: swim, swimming, etc.); gestural units can also be lumped at a higher-level (*Raise*) or specified more finely (*Raise-arm* or *Raise-arm + finger-flexion*) [46–48, 50]. However, these gestural units have been variably described across studies and species with differences in the degree of lumping and splitting even within the same study’s repertoire [46]. Uneven lumping and splitting across a repertoire can create a bias in the assessment of how frequently a particular unit is used (a higher-level unit will be detected more frequently than more finely-split units).

These questions are not trivial aspects of methodology specific to ape gesturing—the same decisions over how to lump or split, and their subsequent systemic biases in units and measurements, are found across comparative studies of other species’ communication [46]. In languages, lumping words into part-of-speech categories or breaking words into letters causes Zipf’s law for word frequencies to disappear [42–43]. There is no universally ‘correct’ level of unit: decisions over which units to measure and how to measure them should be shaped by the question being asked and the data available [46–48]. Where our goal is phylogenetic comparison, we are dependent on comparative analysis across species and families; here, variation in the presence of a linguistic law within a system may itself be a useful feature. The presence or absence of Zipf’s law in different units could serve as an indicator of a question-appropriate level of segmentation for detecting units of information in other species’ communication that can be meaningfully compared to lexical units or word chunks in human languages [43].

The current apparent absence of Zipf’s law in chimpanzee gestural communication may be indicative of a) a genuine system-wide absence in chimpanzee or ape gestural communication, b) a context-specific or population-specific effect, or c) a mismatch between the ways in which efficiency operates on gestural systems and the methods currently used to detect it [6, 32]. Here we test these hypotheses by assessing the presence of Zipf’s law of brevity in the gestural repertoires of three wild chimpanzee communities from two different subspecies. We focus on different levels of segmentation and granularity, namely two different gesture unit durations (Minimum Action Unit duration and Performed Action Unit duration) and two levels of gesture definitons (Gesture Actions and Gesture Morphs). We underscore the importance of considering the social unit and species-specific relevance of units and measures for more meaningful like-with-like comparison across communication system and suggest that socio-ecological dynamics may also influence the expression of Zipf’s law in ape gesturing.

## Results

We provide a multi-level analysis of Zipf’s law in chimpanzee gesture across three communities of two subspecies of chimpanzees (two neighbouring Eastern chimpanzee communitites: Sonso and Waibira, *Pan troglodytes schweinfurthii*; and one Western chimpanzee community: Bossou, *Pan troglodytes verus*), without restricting our analysis to specific contexts or goals (see Supplementary Figure S1, Table S1).

We applied an informed, data driven approach to classifying gesture types uniformly within a repertoire, and at two levels of granularity: Gesture Actions (GAs; lumped) and Gesture Morphs (GMs; split) [47–48]. Gesture Actions were discriminated by their basic movement forms (e.g., *Reach, Big Loud Scratch, Hit*) [47, 48]. Gesture Morphs describe specific consistently produced combinations of Gesture Actions together with a set of modifiers (e.g., limb used: *Reach-arm, Reach-hand*; or repetitiveness: *Hitting*), and were identified by using a Latent Class Analysis (Supplementary Figure S2, Table S2) [48].

We then consider two measures of duration, differentiating the duration of the Minimum Action Unit (MAU) from the duration that includes both the MAU and any hold or repeSSon phase, which we term the Performed Action Unit (PAU; Figure 1). We performed four linear models, one for each combination of the two unit-definitions and two unit-durations, to investigate whether, how, and where compression acts on ape gestural systems, and to investigate the importance of considering signal units and measures relevant to the study species in interspecies comparisons of communication. In addition, given that the expression of Zipf’s law could be influenced by socio-ecological variation between groups, we included a comparison of its expression across three chimpanzee communities.

Specifically, we assessed the effects and interactions of relative Gesture Action frequencies (*rFq_action)* and relative Gesture Morph frequencies (*rFq_morph)* with community (fixed effects) on the MAU or PAU durations of the tokens (response variable), while controlling for signaller identity, goal, and type of Gesture Action or Gesture Morph (included as random effects).

We used video recordings of the Gestural Origins database [51], to measure n=7831 gesture tokens (Sonso = 4515, Waibira = 1423, Bossou = 1893) produced by 175 signallers (Sonso = 72, Waibira = 76, Bossou = 30) across the three chimpanzee communities (Supplementary Section 3). As predicted, the MAU durations showed more conserved durations, with substantially less variability within each gesture action compared to the PAU durations (MAU sd range= 0.07-1.71: ; PAU sd range = 0.07-19.69; Supplementary Figure S3).

All four full models, which assessed the effect of the interaction between frequency of Gesture Actions (*rFq_action*) or Gesture Morphs (*rFq_morph*) and community on the MAU duration of tokens (MAU-Gesture Action and MAU-Gesture Morph models), and on the PAU duration of tokens (PAU-Gesture Action and PAU-Gesture Morph models) performed significantly better than their respective null models (MAU-Gesture Action: X^2^_(5)_=64.69, p<0.001; MAU-Gesture Morph: X^2^_(5)_=69.29, p<0.001; PAU-Gesture Action: X^2^_(5)_=45.28, p<0.001; PAU-Gesture Morph : X^2^_(5)_=69.29, p<0.001). These results suggest that the MAU and PAU durations of chimpanzee gestures vary based on the frequency of Gesture Actions (*rFq_action*) or Gesture Morphs (*rFq_morph*) and/or community. For both MAU models (which had MAU durations as response variables) we retained the interaction factor between *rFq_action* or *rFq_morph* and community as it significantly improved model performance (MAU-Gesture Action: LRT, X^2^=9.85, p=0.007; MAU-Gesture Morph: LRT, X^2^=8.72, p=0.013), highlighting a variation in the effect of *rFq_action* or *rFq_morph* on the MAU duration of Gesture Actions and Gesture Morphs between communities (Supplementary Tables S4 and S8; Figure 1).

For both PAU models the interaction factor did not improve model performance (PAU-Gesture Action: LRT, X^2^=4.83, p=0.09; PAU-Gesture Morph: LRT, X^2^=53.84, p=0.14), suggesting no community variation in the relationship between gesture frequency and PAU duration. When re-computing both PAU models including the relative frequency of the Gesture Actions *rFq_action*, or the relative frequency of the Gesture Morphs *rFq_morph*, and community as separate fixed factors, we found no significant effect of *rFq_action* or *rFq_morph* on PAU duration, suggesting a general lack of Zipf’s law across all communities at the level of PAU durations. The only variation detected was given by community differences in gesture PAU duration (Supplementary Tables S12; S13).

### Community-specific subsets

Given that the relationship between *rFq_action* and *rFq_morph* on the MAU duration of Gesture Actions and Morphs differed across communities, we re-calculated both MAU models for each community separately in order to determine whether Zipf’s law was present in particular communities. We detected no significant relationship between *rFq_action* and Gesture Action MAU duration in either Sonso or Waibira communities (Supplementary Tables S5, S7). However, we found a negative correlation between *rFq_action* and Gesture Action MAU duration within the Bossou community, suggesting the presence of Zipf’s law (Estimate = -6.27 ± 2.95, 95% CI [-12.06; -0.48], p<0.034; Figure 2), although the effect was weak (Marginal R^2^=0.047, Supplementary Table S6). The relative frequency of Gesture Morphs *rFq_morph* did not affect MAU duration in any of the community-specific subsets (Supplementary Tables S9-11).

**Figure 2.**
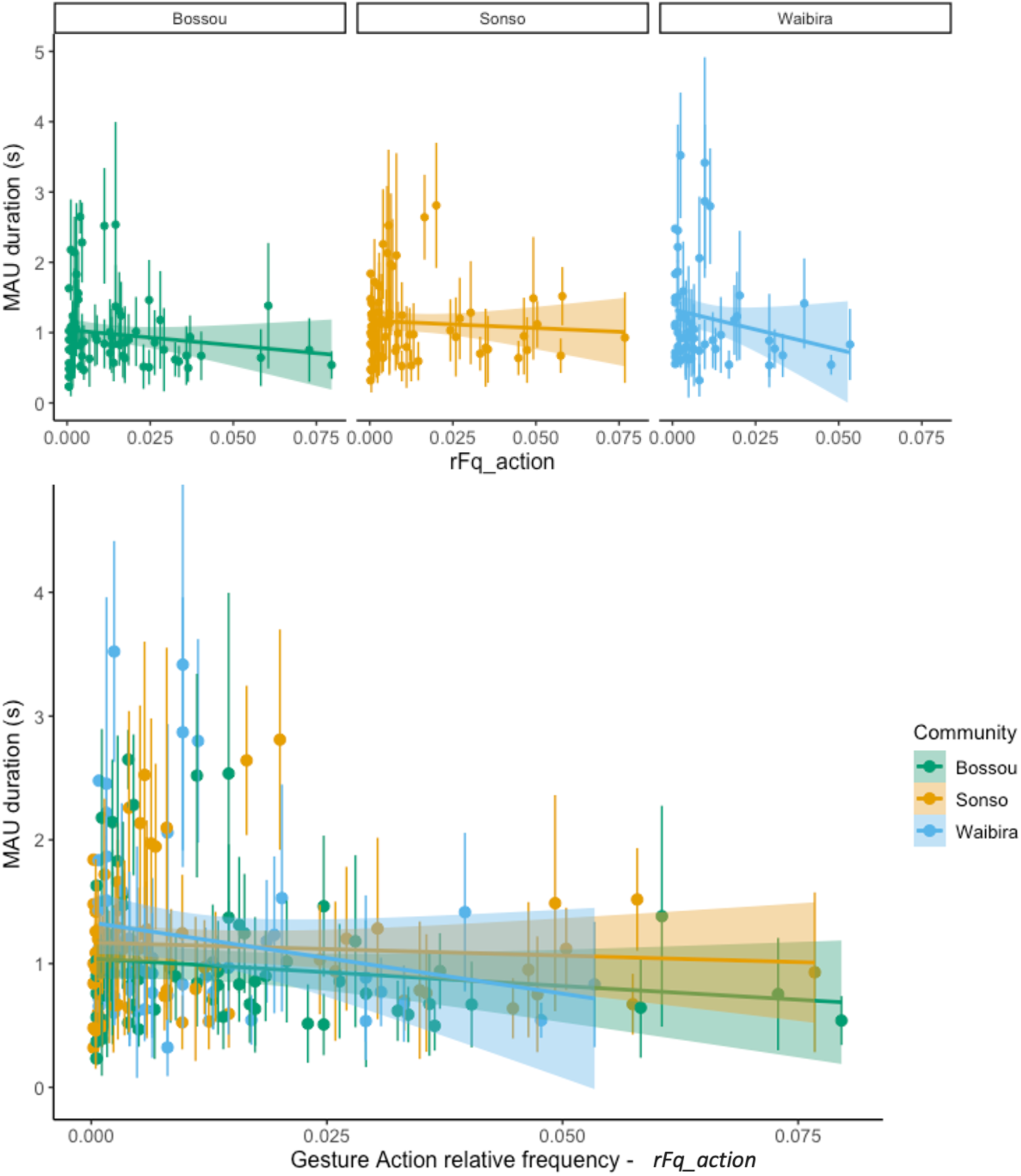
Relationship between the MAU duration of Gesture Actions and their relative frequency (*rFq_action*). Points indicate mean Gesture Action duration, bars indicate standard error, colours indicate community, shaded area indicates 95% confidence intervals of regression line.

## Discussion

We report the detection of Zipf’s law of brevity in the full gestural repertoire of one chimpanzee community when assessed by a specific duration and gesture classification measure. Zipf’s law was present in the Bossou Western chimpanzee community, when assessing the Minimum Action Unit duration of Gesture Actions, suggesting that this is the unit on which compression acts in at least one chimpanzee gestural repertoire. The Minimum Action Unit and Gesture Action level both represent the core components of gesture production, present in every instance of gesture production of this action type. They respectively describe the phases and the form of the gesture that are robust across uses, individual expressions and immediate circumstances. Gesture Actions appear to be shared both across communities of the same species and across species of the same family, therefore being phylogenetically conserved [52]. As a result, this level of segmentation and granularity are likely well suited to detecting selection pressures that act on species-level repertoires of communicative units. In contrast, Performed Action Units (as well as the full duration used in previous studies [31–32]) incorporate optional phases sensitive to the immediate socio-ecological circumstances of each specific communication. Signallers may—case by case—add information (by holding or repeating the action), based on the age, sex, receptiveness, and social status of the recipient and themselves, the nature of their relationship, or the wider social and behavioural context of the event or day [49]. As a result Performed Action Unit durations are highly variable across instances of use.

The absence of any effect of Zipf’s law in the Gesture Morph repertoire in all three communities suggests that compression is more likely to act at the level of the Gesture Action units, as opposed to the more finely split Gesture Morphs. Morphs incorporate information on additional elements of gesture expression (the articulator used, the incorporation of objects, etc.), and could offer a means to reduce energetic investment (for example *Raise* of a single arm, or just the hand, rather than two arms). One alternative possibility, is that Gesture Morphs may instead serve to specify or refine the information transmitted, or may reflect individual or local variation in gesture expression (e.g., [53]).

We remain cautious in our interpretation of any apparent difference in the expression of Zipf’s law between Gesture Morphs and Gesture Actions. Gesture Morphs are currently established by considering patterns of co-occurrence in the use of actions and modifiers that suggest a distinct signal boundary [47]. However, Gesture Morphs are more easily detected from those Gesture Actions that are well-represented in the dataset. With double the number of Gesture Morphs (140) as compared to Gesture Actions (70) there is an impact on data density and any associated statistical power (even in our relatively large dataset for gestural studies). Thus, we remain agnostic as to whether the absence of Zipf’s law at the Gesture Morph level represents a property of the chimpanzee gestural system or limitations in our methods or datasets. At the same time, while our sample size is comparatively small when compared to linguistic corpora (which can include over two-million tokens; [54]), if any absence was primarily a matter of data density, the strongest pattern of compression should emerge in the community with the largest dataset per unit of analysis—in our data, Sonso—and this was not the case.

Our results showcase the importance of assessing diverse measures and units when comparing different species’ systems of communication. The lack of any effect of Zipf’s law on the Performed Action Unit across all groups, suggests that the inclusion of optional elements of gesture performance may mask the expression of this law [31–32]. Zipf’s law of brevity is a system-level linguistic ‘law’, relevant to compression across the information present in all uses of a signal. Duration of the whole gesture unit is not equivalent to the duration of a whole spoken word—we do not continuously ‘hold’ the production of a word or phoneme when waiting for a response. By selecting units of analysis more appropriately tailored to the studied species and communicative channel, we enabled a more direct comparison of like-with-like duration measures and demonstrated a first detection of Zipf’s law in ape gesturing at the whole-repertoire level.

At the same time, we only detected Zipf’s law in the gesturing of a single community. Given its widespread presence across communication systems, its presence remains the default expectation. One possible explanation is that gesture duration—or duration alone—does not represent the most accurate measure of energetic investment in ape gesture, or that it may vary between communities. While duration offers one measure of energetic expenditure in gesture production, others could include the size or speed of movement (similarly to amplitude in vocalizations [12]). Researchers have yet to explore the three-dimensional space of gestural production, but advancements in the automation of pose estimation (e.g., [55]), may allow for these measures—or composite measures— to be considered in the future.

The absence of Zipf’s law patterns in any measure in either Eastern chimpanzee community suggests that the effects of compression may also be sensitive to socio-ecological variation. In human experiments where participants had to learn and use an artificial language, Zipf’s law only emerged in the task where participants faced pressure towards both accuracy and time-efficiency [56], both of which may vary based on the context of communication [31–32] or the social dynamics between signaller and recipient, or of the group as a whole [49].

The differences we see in the expression of Zipf’s law between communities may result from the different socio-ecological pressures faced by individuals within each community. While we found the presence of Zipf’s law of brevity in the small and highly cohesive Bossou community, it was absent from the two larger and more dispersed Eastern communities of Sonso and Waibira. This inter-community variation is congruent with the ways in which Zipf’s law emerges in other systems. For compression of a communicative system to occur there must be repeated instances of use—Zipf’s law depends on the principle that more frequently used units are subject to greater compression. Given the evidence that signaller and recipient identity, as well as behavioural context [57–58], shape gesture meaning, stronger compression may occur where there are repeated instances of gesture use between the same interlocuters towards the same goals, as is more likely to be found in small and cohesive groups. Bossou is a particularly small and isolated community with little-to-no immigration from neighbouring communities (5-30 individuals during the study period) [59], and the individuals form a highly cohesive group (as is more typical for Western chimpanzees [60]), frequently traveling as a single party [61]. In contrast, Sonso and Waibira consist of around 70 and over 120 individuals respectively, with regular immigration and greater reliance on fission-fusion dynamics that leads to lower group cohesion [62]. Gestural communication, like spoken or signed languages, is primarily used in face-to-face interactions [27]. Communities that frequently act as a single cohesive group likely experience greater repetition of gesture use between the same dyads for the same goals, which may favour selection for compression. In line with this hypothesis, future research could focus on dyadic communicative efficiency, assessing whether individuals who exchange gestures more often show a trend towards compression even in communities where Zipf’s law of brevity is absent in the overall gestural repertoire.

Conversely, there may also be social contexts in which gestural communication would benefit from *less* compression, for example where there is a trade-off between accuracy and reducing miscommunication [31–32, 63]. The apparent absence of compression in Sonso and particularly in the ‘super-sized’ Waibira community, which shows additional levels of social structure beyond fission-fusion dynamics [64], could reflect energetic investment in gesture as a marker of costly signalling. Suggested as a precursor to ritualised indications of co-operative intention [65], in human communities, costly signalling tends to increase with increasing group size and may serve to stabilise larger social networks [66]. Given the lower likelihood of encountering some—perhaps many—of the individuals in their community on a regular basis, chimpanzees in Sonso and, especially, Waibira may experience both a higher risk of miscommunication and fewer opportunities for reconciliation; as a result they may favour redundant, repetitive, or longer communicative signals as a risk-mitigation strategy [67].

Human language has been suggested by some to originate in a system akin to modern ape gesture [68–71], supported by the abundant evidence for key linguistic capacities including intentional meaning [26, 72–73] and substantial means-ends flexibility [58, 74]. The apparent absence of Zipf’s law—robustly found across expressions of human language, and widespread in many communication systems—has been puzzling. Given that research has highlighted the presence of Zipf’s law across diverse natural systems [10–16, 23–24], it remained unclear why a system that shares so many of the properties of human language does not appear to reflect the same patterns of efficiency. The application of quantitative linguistics to ape gesture remains a very young field [31–32], and the results of exploratory work are likely shaped by the ways in which our methods are adapted from human studies. Our findings highlight that identifying the appropriate unit of analysis is important for detecting Zipf’s law expression. The detection of linguistic laws in languages is sensitive to similar methodological decisions on the level of granularity or lumpling [42–43, 75]. When approaching other species, researchers lack the knowledge that linguists have as (language-competent) ‘insiders’ in order to make appropriate methodological decisions—whether for the detection of Zipf’s law or other comparative analyses. While different units are likely appropriate for different questions; we suggest that establishing at what level of unit Zipf’s law is detected in a species repertoire may provide a useful indicator of an appropriate level for comparison across linguistic features beyond Zipfian distributions.

## Methods

### Study site & species

We analysed videos of three chimpanzee communities: the Sonso and Waibira communities (Eastern chimpanzees, *Pan troglodytes schweinfurthii*) of the Budongo Forest Reserve, Uganda (1°35ʹ and 1°55ʹ N and 31°08ʹ and 31°42ʹ E), and the Bossou community (Western chimpanzees, *Pan troglodytes verus)* in Guinea (7°39ʹ N and 8°30’W). See Supplementary Section 6 for video collection procedures.

### Video coding

N=2640 videos were coded following the GesturalOrigins protocol [48] using ELAN [76], which allows for accurate duration measurements up to 0.04s. We included both successful and unsuccessful communications, discriminating gestural instances initially on their basic movement forms, termed Gesture Actions (GAs). Each Gesture Action is annotated on a continuous timescale, marking Gesture Action start and end (Supplementary Section 6). We then coded a set of production characteristics (termed *modifiers*), including but not limited to: (1) body part used to perform the gesture action (limb and laterality, e.g., arm right), (2) directionality (e.g., if the gesture was directed to a location) and (3) repetitiveness (e.g., stomp vs stomping; see Supplementary Figure S2, Supplementary Table S2). We applied a Latent Clustering Analysis to describe systematic subcategories of Gesture Actions + modifiers, based on gesture use, to define a repertoire of distinct units we term *Gesture Morphs* (GMs) [47]. In addition, we coded signaller and recipient identities and the apparent communicative goal (see Supplementary Section 2).

### Relative frequencies of Gesture Actions (*rFq_action*) and morphs (*rFq_morph*)

To account for uneven sampling across communities, we calculated ‘Relative frequency’ of types at two levels of granularity: Gesture Actions (*rFq_action*) and Gesture Morphs (*rFq_morph*) by dividing the number instances of a particular type of Gesture Actions (for *rFq_action*) or Gesture Morphs (for *rFq_morph*) within a community, by the total number of gesture instances from that community.

### Inter-observer reliability

Reliability of coding was assessed by conducting ICC, Cohen’s Kappa, or percentage agreement on selected variables, on a subset of the data, representing around 5% of the total gestures coded. Overall agreement between the two coders for the Eastern chimpanzee dataset (ASa, GB) was excellent for both MAU duration (ICC= 1) and PAU duration (ICC=1). There was substantial agreement on the identification of Gesture Actions (Cohen’s Kappa=0.798), and agreement on the modifiers employed in the clustering analysis for Gesture Morph creation was near perfect (Repetition (Y/N) Cohen’s Kappa=0.952; Body part, Cohen’s Kappa=0.856). Agreement on communicative goal was substantial (Cohen’s Kappa=0.786). Overall agreement between the two coders for the Western chimpanzee dataset (ASa, DR) was again excellent for both MAU duration (ICC= 1) and PAU duration (ICC=1). There was also almost perfect agreement on the identification of Gesture Actions (Cohen’s Kappa=0.801), and the other modifiers employed in the clustering analysis for Gesture Morph creation matched substantially (Repetition (Y/N) Cohen’s Kappa=0.699; Body part, Cohen’s Kappa = 0.793). Agreement on communicative goal was almost perfect (Cohen’s Kappa = 0.874). Full details of the IOR procedure can be found in Supplementary Section 7.

### Statistical analyses

Data were analysed using R v. 4.3.0 and RStudio v. 1.2.5042 [77–78]. Zipf’s law was tested on two measures of duration: MAU and PAU, and at two levels of granularity: Gesture Actions and morphs. All measures were tested using general linear mixed models, using the ’lmer’ function from the ’lme4’ package [79]. Given the previous absence of Zipf’s law within one of the study communities (Sonso-Budongo: [31–32]), and the potential for variation in Zipf’s law expression given the variation in gesture use across groups [64], we first assessed whether the relationship between gesture frequency and duration varies between communities and, where relevant, assessed Zipf’s law of brevity in community-specific subsets.

Four models were tested. The MAU-Gesture Action and MAU-Gesture Morph models both contained the MAU duration of tokens as response variable, with MAU-Gesture Action including the relative frequency of Gesture Actions (*rFq_action*), the community, and the interaction between *rFq_action* and community as fixed effects; while MAU-Gesture Morph included the relative frequency of Gesture Morphs (*rFq_morphs*), the community, and the interaction between *rFq_morph* and community as fixed effects. PAU-Gesture Action and PAU-Gesture Morph were assigned the same fixed effects, but with the PAU durations of tokens as the response variable. We included signaller identity, communicative goal, and Gesture Action (for -GA models) or Gesture Morph (for -GM models) as random effects. All response variables were log-transformed. All full models were tested against null models via Likelihood Ratio Tests, using the ’anova’ function from the ’stats’ package (version 4.1.2, [77]). The importance of the interaction factor was assessed by performing sequential model comparison using ’drop1’ from the ’stats’ package (version 4.1.2, [77]). Where the interaction factor did not significantly improve the model performance, we removed the interaction and retained each variable as a separate fixed effect. All models were assessed for multicollinearity of factors using the ’vif’ function from the ’car’ package (version 3.1-2, [80]), with variance inflation factors (VIF) for all models being <1.09 [81]. All models were evaluated on the assumptions of linear models by visual inspection of residual distributions (see Supplementary Methods 6). Significance of fixed effects was determined based on assessment of p-values (alpha level = 0.05) and 95% confidence intervals.

## Author contributions

Conceptualisation: A.Sa., and C.H; Methodology: A.Sa., C.G., C.H., and R.F.C; Formal Analysis: A.Sa.; Data Collection: A.Sa., A.So., C.H., C.Z., E.D.R., G.B., and M.H.; Data Curation: A.Sa., E.D.R, and G.B.; Writing original draft: A.Sa.; Visualisation: A.Sa., Writing—reviewing and editing: A.So., C.H., C.G., C.Z., E.D.R., R.F.C., G.B., J.M., M.H. and K.Z.; Resources: C.H., K.Z., and M.H.; Supervision: C.H.; Funding acquisition: C.H., K.Z., and M.H.

## Acknowledgements and Funding

ASa, GB, CG, and CH were supported by funding from the European Research Council under the Gestural Origins Grant No: 802719. ASa was supported by the Japan Society for the Promotion of Science (JSPS, fellowship No: PE22721) for working on the Bossou Video Archive. EDR was supported by national funds from FCT – Fundação para a Ciência e Tecnologia, I.P. (SFRH/BD/138406/2018). RFC is supported by a recognition 2021SGR-Cat (01266 LQMC) from AGAUR (Generalitat de Catalunya) and the grant AGRUPS-2024 from Universitat Politècnica de Catalunya. MH was supported by JSPS KAKENHI 21K18138 and Innovative Areas no. 4903 (Evolinguistics) JP17H06381, and research fund of Chubu Gakuin University.

We thank all the staff of Budongo Conservation Field Station (BCFS) and are grateful to the BCFS founder Vernon Reynolds, and the Royal Zoological Society of Scotland who provide core funding. We thank the Uganda Wildlife Authority, the National Forestry Authority, the President’s Office, and the Uganda National Council for Science and Technology for providing research permits and permissions to conduct research in Budongo. Original Bossou videos were collected by members of KUPRI-International led by Tetsuro Matsuzawa, Dora Biro, and Susana Carvalho primarily funded by MEXT-JSPS #16002001, #20002001, #16H06283, JSPS-HOPE, gCOE-A06-D07, LGP-U04, and CCSN. We are also grateful to Primate Research Institute, Kyoto University for leading the Bossou Archive Project and supporting the video digitization mainly led by Daniel Schofield and Kirsty Graham that enabled the comparative research presented here and to the IREB and DNRSIT of Guinea for giving research permission. This study is dedicated to all the researchers and field assistants who have collected data in Bossou since 1988.

