## Supporting Material for "Testing methodological and population-specific influences on the detection of Zipf’s law of brevity in chimpanzee gestures"

**Supplementary Material – Methods & Results**

1. **Goal distribution**

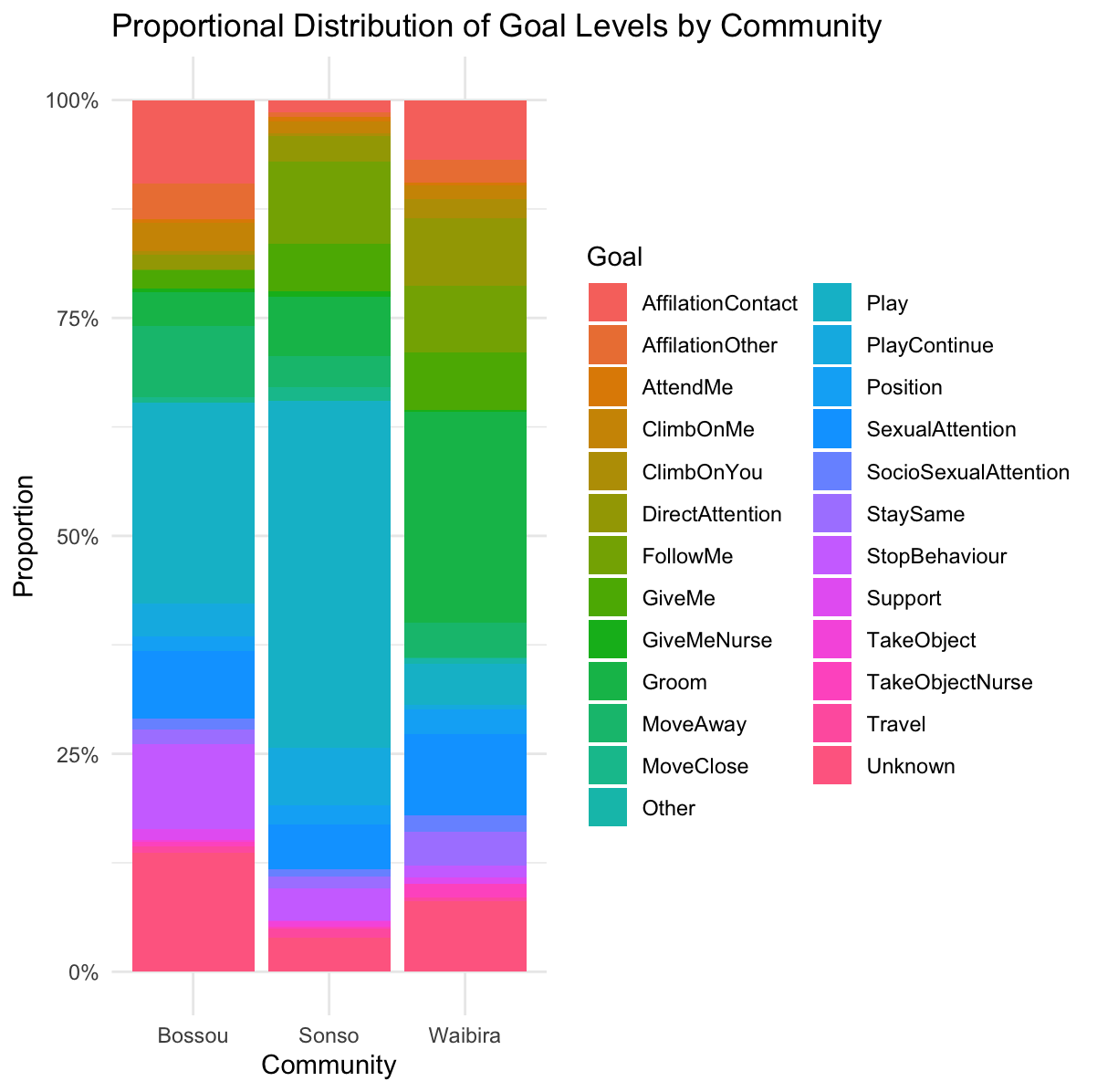

**Figure S1. Distribution of communicative goals across the three communities within the dataset.** For definitions and frequencies please consult Table S1 below.

Table S1. Goal definitions and lumping. Adapted from [1] and [2]. Goal lumping indicates grouping made to decrease variability of this variable used as random effect in multiple models. Frequencies of each lumping group are reported per community.

| **Original coding** | **Definition** | **Lumping** | **Bossou** | **Sonso** | **Waibira** |
| --- | --- | --- | --- | --- | --- |
| Affiliation: Contact | Affiliation with body contact (initiation of affiliation behaviour can be by signaller or recipient). Note that body contact that includes touching of genitals should be coded as socio-sexual. | Affiliation: Contact | 185 | 65 | 105 |
| Affiliation: Rest in contact | Signaller and recipient rest while making physical contact after a gesture(s). |  |  |  |  |
| Affiliation: Distant | Affiliation without body contact (initiation of affiliation behaviour can be by signaller or recipient). | Affiliation: Other | 82 | 23 | 38 |
| Affiliation: Unclear | Affiliation but whether or not body contact occurs is unclear (initiation of affiliation behaviour can be by signaller or recipient). |  |  |  |  |
| Attend Me | Signaller requests that the recipient change their behaviour in order to visually monitor the behaviour of the signaller. Note that this is rarely ever an end goal for non-human apes but could be used if there was a gesture with a theoretical 'attention getter' only function. | Attend Me | 8 | 23 | 4 |
| Climb On Me | Recipient moves (climbs) onto the signaller, usually infant climbs onto mother or another individual. | Climb On Me | 61 | 68 | 20 |
| Climb On You | Recipient allows signaller to climb onto them, usually infant is the signaller and they climb onto mother or other individual. | Climb On You | 7 | 15 | 31 |
| Direct Attention | Normally used during social interactions (e.g., grooming, nursing) the recipient changes the location of this behaviour by focusing their attention on a different body part (e.g., grooms new body part, nurse from the other breast). The recipient must be actively engaged in the social behaviour before the signaller gestures and the recipient reacts by directing the same behaviour somewhere else. | Direct Attention | 30 | 130 | 104 |
| Follow Me | Signaller requests that the recipient follow them in order to perform another activity in another location, for example move over here to groom, have sex, etc. Note that 1) the signaller must gesture before moving to the new area where the behaviour happens. 2) the signaller must be satisfied (i.e. no more response waiting) by the 'following', if they continue to wait until - for example - grooming starts then the outcome that satisfies them would be grooming (not following). Note that this is not about 'travelling together' over longer distances, and also that 'Follow me' as a goal will often not be coded because the final behaviour is often something else (so it gets 'lost’ but can be checked for later in combinations). | Follow Me | 2 | 425 | 99 |
| Give Me | Object/food is exchanged from the recipient to the signaller but it is unclear who takes the active role in the exchange. | Give Me | 39 | 260 | 89 |
| Give Me: Active | The signaller makes an active action to share food with the recipient, for example putting the food in their hands, passing the food to them, extending it towards them so that they can more easily take it, moving in order to drop it where the recipient can access it. |  |  |  |  |
| Give Me: Passive | The signaller permits the recipient to access the food but does not take any overt action to share it. For example, they allow the recipient to approach and pick up food in their hands or mouth, but also food that is near them but within reach. Includes 'tolerated pilfering'. |  |  |  |  |
| Give Me: Nurse | Recipient allows signaller to nurse after the signaller gesture toward recipient. | Give Me: Nurse | 7 | 25 | 3 |
| Groom | Recipient and signaller start grooming together but unclear if the goal was to groom or be groomed. Note that if grooming is ongoing and a gestural request occurs consider whether or not this is a new grooming request, or a request to Direct (grooming) Attention to a new location. | Groom | 72 | 310 | 344 |
| Groom: Me | Recipient starts grooming the signaller. |  |  |  |  |
| Groom: You | Recipient allows signaller to start grooming them. |  |  |  |  |
| Move Away | Recipient typically stops current behaviour and increases physical distance between themselves and signaller. Move away will typically be used when the signaller is stationary. Move away must include more than leaning away and more than 2-steps back (otherwise see: Stop that). | Move Away | 154 | 152 | 55 |
| Move Close | Physical distance between signaller and recipient decreases, normally because recipient moves closer. | Move Close | 12 | 74 | 1 |
| Other | Other goal (not listed). | Other | 0 | 0 | 7 |
| Play: Start | Signaller is resting, engage in a non-play activity, or playing with a different individual and gestures to recipient to initiate play. Signaller may have been playing a short while before with the recipient but they must have changed their activity to another one (feeding, clearly resting without waiting etc.), if not consider Play-continue. | Play | 434 | 1783 | 67 |
| Play: Change | Signaller gestures to recipient and the play-type changes (change unspecified). | Play: Continue | 68 | 289 | 6 |
| Play: Change chase to contact | Signaller gestures to recipient and the play-type changes from non-contact (e.g., chase) to contact (e.g., wrestle). |  |  |  |  |
| Play: Change contact to chase | Signaller gestures to recipient and the play-type changes from contact (e.g., wrestle) to non-contact (e.g., chase). |  |  |  |  |
| Play: Continue | Signaller has been playing with recipient, but play was interrupted by a third party, or there was a short break in which they continue to monitor each other. Signaller gestures in order to resume the play. |  |  |  |  |
| Position | Signaller is usually actively engaged in a social behaviour with the recipient (e.g., signaller grooms recipient/ signaller nurses from recipient). The signaller gestures for the recipient to change their position so that the signaller can continue their behaviour in a different location (i.e., groom the recipient at a different spot, nurse from the other nipple). | Position | 33 | 96 | 42 |
| Sexual Attention | Behaviour linked to reproductive sexual activities (copulation, inspection, etc.). Note that this can be challenging to differentiate from affiliative and socio-sexual behaviour across some species (e.g., gorillas). In chimpanzees and bonobos sexual attention = penetration when female is in swelling, inspection (sniffing, tasting, poking) of female genitals at any stage by a male individual who is older than an infant. | Sexual Attention | 143 | 241 | 138 |
| Sexual Attention: To me | Sexual attention to me (signaller). |  |  |  |  |
| Sexual Attention: To you | Sexual attention to you (recipient). |  |  |  |  |
| Sociosexual Attention | Behaviour that involves genital contact that is not linked to reproductive activities. For example, touching testicles, GG-rubbing etc. Note that socio-sexual behaviour is not necessarily positive (i.e., affiliative) and can involve dominance and other aspects. In chimpanzees and bonobos socio-sexual = all same sex interactions, opposite sex interactions where there are none of the sexual behaviours described above. | Socio-sexual Attention | 24 | 42 | 24 |
| Sociosexual Attention: To me | Socio-sexual attention to me (signaller). |  |  |  |  |
| Sociosexual Attention: To you | Socio-sexual attention to me (recipient). |  |  |  |  |
| Stay Same | Essentially there is no change in the recipient's behaviour but the signaller behaves as if they are satisfied with this outcome. This is a difficult one because it's a 'goal' with no visible change in behaviour. So rather than be dependent on the recipient, look to what the signaller does - do they seem satisfied (no persistence, extended response waiting) by the lack of response. | Stay Same | 31 | 65 | 64 |
| Stop Behaviour | The recipient stops the behaviour they were in before the gesture and doesn't change their behaviour to something that we have as a possible goal the signaller might be asking for. This typically means that they stop all activity and pause or rest for a (brief) time, often seen in contexts such as begging, but can be used across contexts. | Stop Behaviour | 182 | 160 | 21 |
| Stop Behaviour: Stay | Signaller and recipient are near to each other, the recipient stands up as if to move away, the signaller gestures and the recipient does not leave but stays close. |  |  |  |  |
| Stop Behaviour: Here | Recipient's behaviour does not completely stop but is continued at a different place. E.g., signaller and recipient are feeding on bark with heads close. Signaller gestures to Recipient and recipient moves a bit away and continues feeding at a different spot (so behaviour of recipient is the same as before the gesture but at a different spot - which seems to sufficiently satisfy the signaller). |  |  |  |  |
| Support | After the gestural interaction signaller and recipient direct behaviour - typically aggressive/agonistic - toward another individual. Aggression is one possible outcome, but other outcomes would be vocal support (bark), and the support should include behaviour that is directed to a third party. If positive behaviour is directed to the recipient see affiliation goals. Unclear who is giving who the support. | Support | 24 | 9 | 11 |
| Support: Me | See Support but clear that recipient is the one to give subsequent support to signaller. |  |  |  |  |
| Support: You | See Support but clear that signaller is the one to give subsequent support to recipient. |  |  |  |  |
| Take Object | Object/food is exchanged from the signaller to the recipient but it is unclear who takes the active role in the exchange | Take Object | 5 | 26 | 0 |
| Take Object: Active | The signaller makes an active action to share food with the recipient, for example putting the food in their hands, passing the food to them, extending it towards them so that they can more easily take it, moving in order to drop it where the recipient can access it. |  |  |  |  |
| Take Object: Passive | The signaller permits the recipient to access the food, but does not take any overt action to share it. For example, they allow the recipient to approach and pick up food in their hands or mouth, but also food that is near them but within reach. Includes 'tolerated pilfering'. |  |  |  |  |
| Take Object: Nurse | Recipient nurses from signaller, after the signaller gesture toward recipient. | Take Object: Nurse | 9 | 9 | 21 |
| Travel | Signaller gestures to another individual in the party, and response waits or checks back to recipient. Both travel in the same direction and within sight. | Travel | 15 | 46 | 6 |
| Travel: With me | Signaller gestures to another individual in the party, and response waits or checks back to recipient until the recipient travels in the same direction as them and within sight of them. If signaller is already travelling, need evidence that they wait for recipient to also travel with them. If recipient is already travelling, need evidence that they changed their travel to move with the signaller (e.g., direction). Note - if there's no evidence that the signaller waits for the recipient or monitors them then this could not be coded as the goal. The goal would be unknown. |  |  |  |  |
| Travel: With you | Rare. Signaller gestures to another individual in the party, and response waits or checks back to recipient until the recipient pauses to allow recipient to travel in the same direction as them and within sight of them. Note - if there's no evidence that the signaller waits for the recipient or monitors them then this could not be coded as the goal. The goal would be unknown. |  |  |  |  |
| Unknown | Unknown which goal the signaller wanted to achieve. | Unknown | 266 | 179 | 123 |

2. **Gesture Morphs**

So far, in gestural research, studies have followed a diverse set of gesture definitions for communicative units, usually based on a mixture of morphological and contextual definitions designed to achieve high reliability within studies, but that offered limited consistency between studies [1, 3]. The application of pre-defined repertoires has the potential to divide gesture units at arbitrary points at which characteristics that appear important to a human coder may be irrelevant for the individuals employing these gestures, or vice versa where details relevant to the chimpanzees are left uncoded by researchers.

For instance, chimpanzees' contact gestures that involve a similar action movement but with different body parts: fingers (*Poke*), palm (*Slap*), or fist (*Punch*), exhibited very similar goal-oriented patterns [5]. Thus, while human observers might sometimes categorize them separately as distinct 'types' [6], it is less clear that the chimpanzees using them would do the same. It is important to describe a signal repertoire that allows researchers to detect and define units that reflect relevant granularity for the user (here chimpanzees), rather than the coder. As well as some flexibility to lump or split these depending on the data available and the scientific question of interest. With the Gestural Origins protocol [1], we wanted to find a standard coding scheme that allowed for the encoding of basic gestural action and then creation of distinct gestural repertoires depending on the systematic application of a set of characteristics, previously named *features* in other publications [e.g., 2, 5]. In constructing our repertoire from the bottom-up we have the opportunity to re-construct gesture units flexibly in ways that best reflect species-specific use, and suitability for analysis. We term the gesture units created by the addition of modifiers to Gesture Actions, gesture Morphs. By performing a Latent Class clustering analysis, we discriminated which combination of Gesture Action plus modifier appeared to be consistently distinguished from other Gesture Action plus modifier combinations within the dataset, from the perspective of the chimpanzees using them [2]. The clustering analysis resulted in the detection of 140 distinct gesture Morphs, which represent communicative gesture units at a higher level of *granularity*. Each Morph is characterised by a set of *rules* that describe the combination of modifier levels that led to the creating of that Morph (Figure S1; Table S1). The degree of specificity of gesture Morphs depends on the modifiers included within the clustering analysis, as well as the available data: the use of Gesture Actions is entirely described by a single gesture Morph as some Gesture Actions are performed in the same way across almost all uses (e.g., *Dangling* can only be performed by the whole body as by definition this Gesture Action is limited to this body part). Other Gesture Actions may be split into a number of Morphs (e.g., hitting with one limb or two may result in separate clusters; [2]). It is worth noting that as in linguistic analyses, where phonemes, syllables, and words are all valid levels of examination, Gesture Actions, Morphs, and even more finely divided units are all legitimate levels of gesture analysis – and that researchers can select from within them depending on the research question being asked.

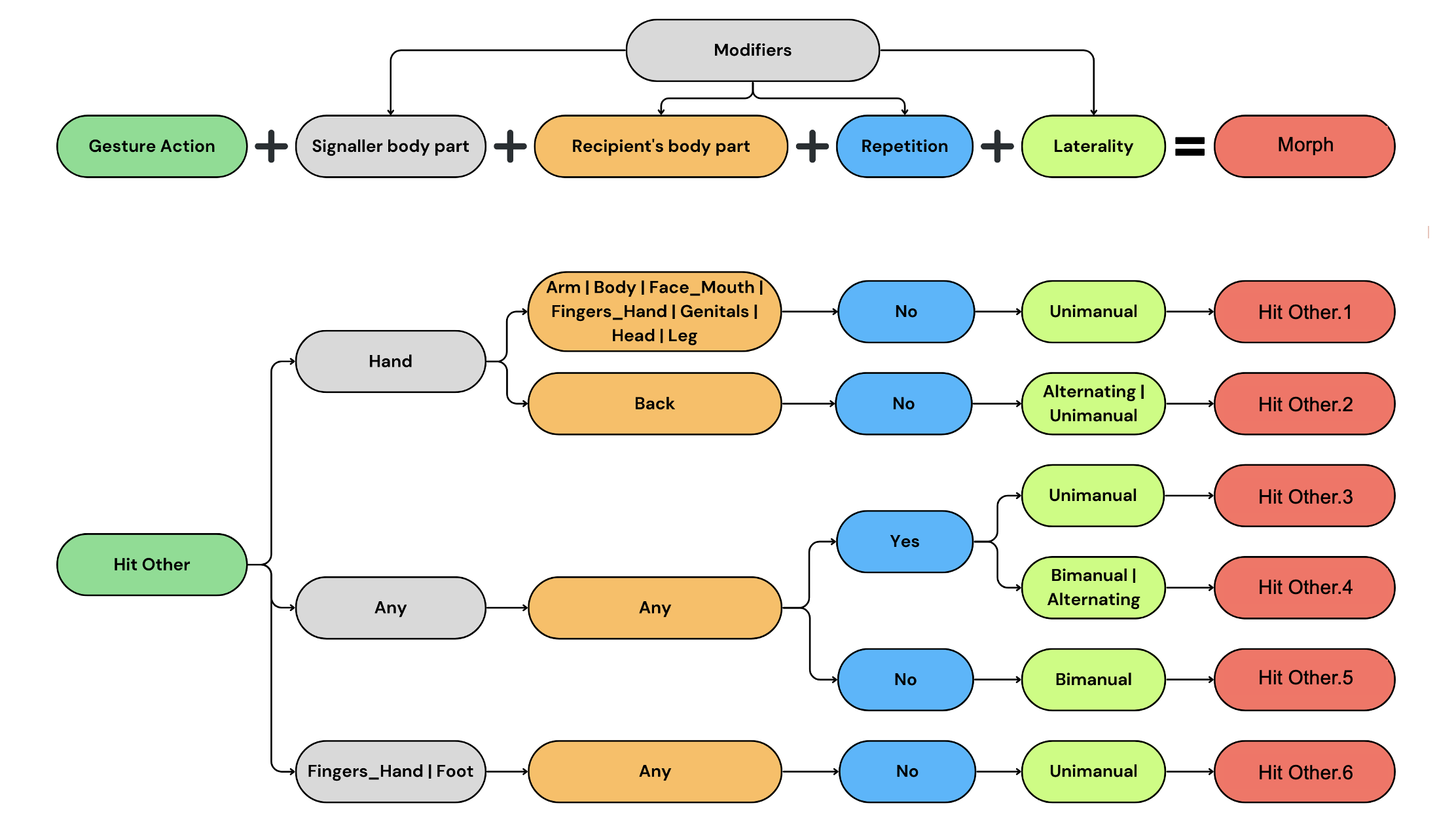

**Figure S2. Flow diagram that explains the creation of Morphs from Gesture Actions and modifier combinations**. Here we report the Morph set derived from the *Hit Other* Gesture Action and modifier combinations. | indicates “or”.

Table S2. List of Gesture Action and modifier combinations with the respective Morphs created via clustering analysis. Modifiers include Signaller’s body part, Recipient’s body part – if contact was involved, whether the gesture was repetitive (Repetition: yes/no), and whether it was performed with one, both or alternating limbs (Laterality: unimanual / bimanual / alternating). | indicates “or”. Frequency of each Morph in each community is also reported. In grey are highlighted Morphs that are not present in this dataset but that were detected in the clustering analysis that contained data from other East African communities (Kalinzu, Kanyawara, Issa).

| **Gesture Action** | **Signaller Body Part** | **Recipient Body part** | **Repetition** | **Laterality** | **Morph** | **Bossou** | **Sonso** | **Waibira** |
| --- | --- | --- | --- | --- | --- | --- | --- | --- |
| Beckon | Arm | NA | NA | NA | Beckon.1 | 18 | 1 | 3 |
| Beckon | Hand | NA | NA | NA | Beckon.2 | 5 | 6 | 5 |
| BigLoudScratch | NA | Body | NA | Unimanual | BigLoudScratch.1 | 6 | 97 | 129 |
| BigLoudScratch | NA | Arm | NA | Unimanual | BigLoudScratch.2 | 28 | 87 | 60 |
| BigLoudScratch | NA | Head | NA | Unimanual | BigLoudScratch.3 | 2 | 54 | 59 |
| BigLoudScratch | NA | Leg | NA | NA | BigLoudScratch.4 | 6 | 19 | 28 |
| BigLoudScratch | NA | Back \| Face_Mouth | NA | Unimanual | BigLoudScratch.5 | 3 | 3 | 9 |
| BigLoudScratch | NA | NA | NA | Bimanual | BigLoudScratch.6 | - | - | 4 |
| Bite | NA | Back | NA | NA | Bite.1 | 2 | 20 | 8 |
| Bite | NA | None | NA | NA | Bite.2 | 24 | - | - |
| Bite | NA | Body | NA | NA | Bite.3 | 4 | 14 | 3 |
| Bite | NA | Arm | NA | NA | Bite.4 | 4 | 8 | 4 |
| Bite | NA | Face_Mouth | NA | NA | Bite.5 | 14 | 5 | 3 |
| Bite | NA | Head | NA | NA | Bite.6 | 4 | 12 | 1 |
| Bite | NA | Fingers_Hand | NA | NA | Bite.7 | 7 | 6 | 3 |
| Bite | NA | Leg | NA | NA | Bite.8 | 1 | 13 | 2 |
| Bounce | NA | NA | NA | NA | Bounce.1 | 38 | - | 1 |
| Bow | NA | NA | NA | NA | Bow.1 | 1 | 8 | - |
| Dangle | NA | NA | NA | NA | Dangle.1 | 48 | 218 | 1 |
| DangleShake | NA | NA | NA | NA | DangleShake.1 | 5 | 32 | 2 |
| Drum | NA | NA | NA | NA | Drum.1 | - | 15 | - |
| Embrace | NA | Body | NA | Bimanual | Embrace.1 | 3 | 13 | 10 |
| Embrace | NA | Body | NA | Unimanual | Embrace.2 | 5 | 9 | 5 |
| Embrace | NA | Back | NA | Unimanual | Embrace.3 | 4 | 1 | 6 |
| Embrace | NA | Back | NA | Bimanual | Embrace.4 | - | 1 | 2 |
| FingerMouth | NA | NA | NA | NA | FingerMouth.1 | 8 | - | - |
| Fling | Arm | NA | NA | NA | Fling.1 | 105 | 31 | 23 |
| Fling | Hand | NA | NA | NA | Fling.2 | 39 | 22 | - |
| Grab | NA | Body \| Face_Mouth \| Head | NA | Unimanual | Grab.1 | 7 | 37 | 5 |
| Grab | NA | Leg | NA | Unimanual | Grab.2 | 8 | 36 | 1 |
| Grab | Hand | Arm | NA | Unimanual | Grab.3 | 15 | 21 | 1 |
| Grab | NA | Back | NA | Unimanual | Grab.4 | - | 34 | 4 |
| Grab | NA | NA | NA | NA | Grab.5 | 10 | 15 | 4 |
| Grab | NA | NA | NA | Bimanual | Grab.6 | 3 | 16 | - |
| GrabHold | NA | Arm \| Back \| Body \| Face_Mouth \| \|Fingers_Hand \| Head | NA | Unimanual | GrabHold.1 | 4 | 42 | 8 |
| GrabHold | NA | Leg | NA | Unimanual | GrabHold.2 | 3 | 25 | 5 |
| GrabHold | NA | NA | NA | Bimanual | GrabHold.3 | - | 24 | 1 |
| HeadStand | NA | NA | NA | NA | HeadStand.1 | 3 | 30 | 1 |
| HitObject | Hand | NA | no | Unimanual | HitObject.1 | 61 | 194 | 58 |
| HitObject | Hand | NA | no | Bimanual | HitObject.2 | 9 | 54 | 5 |
| HitObject | Foot | NA | no | NA | HitObject.3 | 10 | 23 | 6 |
| HitObject | NA | NA | NA | NA | HitObject.4 | 11 | 27 | 6 |
| HitObject* | Hand | NA | yes | Unimanual | HitObject.5 | 10 | 20 | 5 |
| HitObject* | Foot | NA | yes | NA | HitObject.6 | 3 | 7 | 3 |
| HitOther | Hand | Arm \| Body \| Face_Mouth \| Fingers_Hand \| Genitals \| Head \| Leg | no | Unimanual | HitOther.1 | 49 | 91 | 9 |
| HitOther | Hand | Back | no | Alternating \| Unimanual | HitOther.2 | 15 | 42 | 2 |
| HitOther* | NA | NA | yes | Unimanual | HitOther.3 | 1 | 38 | 10 |
| HitOther* | NA | NA | yes | Alternating \| Bimanual | HitOther.4 | 8 | 33 | - |
| HitOther | NA | NA | no | Bimanual | HitOther.5 | 5 | 27 | 1 |
| HitOther | Fingers_Hand \| Foot | NA | no | Unimanual | HitOther.6 | 1 | 22 | 5 |
| Jump | NA | NA | NA | NA | Jump.1 | 24 | 33 | - |
| KickPunch | NA | NA | NA | NA | KickPunch.1 | - | 10 | - |
| KnockObject | NA | NA | NA | NA | KnockObject.1 | 3 | 6 | - |
| LeafClip | Hand | NA | NA | Unimanual | LeafClip.1 | - | 20 | 4 |
| LeafClip | Face_Mouth | NA | NA | NV | LeafClip.2 | - | 6 | 12 |
| LocomoteBipedal | NA | NA | no | NA | LocomoteBipedal.1 | 21 | - | 2 |
| LocomoteBipedal | NA | NA | yes | NA | LocomoteBipedal.2 | 8 | - | - |
| LocomoteGallop | NA | NA | NA | NA | LocomoteGallop.1 | 25 | 37 | 1 |
| Lunge | NA | NA | NA | NA | Lunge.1 | 4 | 2 | 7 |
| ObjectMouth | NA | NA | NA | NA | ObjectMouth.1 | 1 | 21 | 2 |
| ObjectMove | Hand | NA | NA | Unimanual | ObjectMove.1 | 36 | 143 | 26 |
| ObjectMove | Hand | NA | NA | Bimanual | ObjectMove.2 | 23 | 29 | 1 |
| ObjectMove | NA | NA | NA | Alternating | ObjectMove.3 | 4 | 5 | - |
| ObjectMove | Foot | NA | NA | NA | ObjectMove.4 | 3 | 3 | - |
| ObjectShake | Hand | NA | yes | Unimanual | ObjectShake.1 | 18 | 336 | 22 |
| ObjectShake | Hand | NA | yes | Bimanual | ObjectShake.2 | 6 | 107 | 6 |
| ObjectShake | Foot | NA | yes | NA | ObjectShake.3 | 1 | 10 | - |
| ObjectShake | NA | NA | no | NA | ObjectShake.4 | 1 | 13 | - |
| ObjectShake | Hand | NA | yes | Alternating | ObjectShake.5 | 1 | 5 | - |
| Out | NA | NA | NA | Unimanual | Out.1 | 19 | - | 3 |
| Out | NA | NA | NA | Bimanual | Out.2 | 8 | - | - |
| OverStance | Arm | NA | NA | Unimanual | OverStance.1 | 13 | - | - |
| OverStance | Body | NA | NA | NV | OverStance.2 | 9 | - | - |
| Poke | NA | NA | NA | NA | Poke.1 | 2 | 12 | - |
| Present | Back | NA | NA | NA | Present.1 | 28 | 77 | 46 |
| Present | Arm | NA | NA | NA | Present.2 | 11 | 61 | 35 |
| Present | Body | NA | NA | NV | Present.3 | 31 | 21 | 58 |
| Present | Leg | NA | NA | NA | Present.4 | 4 | 13 | 40 |
| Present | Bottom | NA | NA | NV | Present.5 | 25 | 31 | 5 |
| Present | Genitals \| Head | NA | NA | NA | Present.6 | 6 | 7 | 8 |
| Present | Body \| Bottom \| Foot | NA | NA | Unimanual | Present.7 | 4 | 11 | 1 |
| PresentGenitals | Genitals | NA | NA | NA | PresentGenitals.1 | 66 | 133 | 66 |
| PresentGenitals | Bottom | NA | NA | NA | PresentGenitals.2 | 4 | 26 | 15 |
| Pull | NA | Back \| Body \| Face_Mouth \| Fingers_Hand \| Other | NA | Unimanual | Pull.1 | 7 | 37 | 11 |
| Pull | NA | Leg | NA | Unimanual | Pull.2 | 6 | 29 | 16 |
| Pull | NA | Arm | NA | Unimanual | Pull.3 | 13 | 23 | 7 |
| Pull | NA | NA | NA | Bimanual | Pull.4 | 3 | 24 | 3 |
| Push | Hand | Back | NA | Unimanual | Push.1 | 24 | 48 | 16 |
| Push | Hand | Leg | NA | Unimanual | Push.2 | 10 | 16 | 16 |
| Push | Hand | Arm | NA | Unimanual | Push.3 | 25 | 23 | 6 |
| Push | Hand | Body \| Face_Mouth \| Fingers_Hand | NA | Unimanual | Push.4 | 15 | 42 | 2 |
| Push | Fingers_Hand | NA | NA | NA | Push.5 | 12 | 30 | 6 |
| Push | Hand | Head | NA | NA | Push.6 | 15 | 30 | 2 |
| Push | NA | NA | NA | Bimanual | Push.7 | 2 | 13 | - |
| Push | Foot | NA | NA | NA | Push.8 | 2 | 6 | 2 |
| Raise | Arm | NA | NA | Unimanual | Raise.1 | 50 | 39 | 36 |
| Raise | Hand | NA | NA | NA | Raise.2 | 1 | 2 | 8 |
| Raise | NA | NA | NA | Bimanual | Raise.3 | 11 | 8 | - |
| RakeSelf | NA | NA | NA | NA | RakeSelf.1 | 10 | - | - |
| Reach | Arm | NA | NA | NA | Reach.1 | 81 | 184 | 115 |
| Reach | Hand | NA | NA | NA | Reach.2 | 51 | 167 | 12 |
| Reach | Foot \| Leg | NA | NA | NA | Reach.3 | 2 | 10 | 8 |
| Rocking | NA | NA | NA | NA | Rocking.1 | 6 | 17 | 12 |
| RollOver | NA | NA | NA | NA | RollOver.1 | 8 | 28 | 1 |
| Rub | NA | NA | no | NA | Rub.1 | - | 20 | - |
| Rub | Genitals | NA | yes | NA | Rub.2 | - | - | - |
| Rub | Bottom | NA | yes | NA | Rub.3 | 3 | 2 | 2 |
| Shake | NA | NA | no | NV | Shake.1 | 26 | 16 | - |
| Shake | NA | NA | NA | Unimanual | Shake.2 | 4 | 21 | 6 |
| Shake | Head | NA | yes | NV | Shake.3 | 33 | - | 1 |
| Shake | NA | NA | NA | Alternating \| Bimanual | Shake.4 | 1 | 18 | - |
| SpinPirouette | NA | NA | NA | NA | SpinPirouette.1 | 6 | 6 | - |
| SpinRoulade | NA | NA | NA | NA | SpinRoulade.1 | - | 22 | - |
| SpinSomersault | NA | NA | NA | NA | SpinSomersault.1 | 2 | 36 | - |
| StanceBipedal | NA | NA | NA | NA | StanceBipedal.1 | 32 | - | 5 |
| StompObject | Foot | NA | no | Unimanual | StompObject.1 | 28 | 152 | 29 |
| StompObject* | NA | NA | yes | Alternating | StompObject.2 | 17 | 67 | 1 |
| StompObject* | NA | NA | yes | Unimanual | StompObject.3 | 8 | 31 | 4 |
| StompObject | Foot | NA | no | Bimanual | StompObject.4 | 8 | 25 | 1 |
| StompObject | NA | NA | no | Alternating | StompObject.5 | 6 | 7 | 6 |
| StompObject | Hand | NA | NA | NA | StompObject.6 | 2 | 6 | - |
| StompObject* | NA | NA | yes | Bimanual | StompObject.7 | 3 | 3 | - |
| StompOther | NA | Back | no | NA | StompOther.1 | 1 | 11 | - |
| StompOther* | NA | NA | yes | NA | StompOther.2 | - | 11 | - |
| StompOther | NA | Other | no | NA | StompOther.3 | - | - | - |
| Stroke | Hand | NA | no | NA | Stroke.1 | - | 5 | 2 |
| Stroke | NA | NA | yes | NA | Stroke.2 | - | 7 | 4 |
| Stroke | Fingers_Hand | NA | no | NA | Stroke.3 | - | 5 | 1 |
| Swing | Arm | NA | NA | Unimanual | Swing.1 | 57 | 193 | 23 |
| Swing | Leg | NA | NA | Unimanual | Swing.2 | 7 | 14 | - |
| Swing | NA | NA | NA | Bimanual | Swing.3 | 4 | 5 | - |
| ThrowObject | NA | NA | NA | NA | ThrowObject.1 | 30 | 11 | 7 |
| ThrowThreat | NA | NA | NA | NA | ThrowThreat.1 | 36 | 1 | - |
| Touch | Fingers_Hand | Arm \| Back \| Body \| Fingers_Hand \| Genitals \| Head \| Leg | NA | NA | Touch.1 | 16 | 47 | 25 |
| Touch | Hand | Back | NA | NA | Touch.2 | 15 | 34 | 12 |
| Touch | Hand | Leg | NA | NA | Touch.3 | 13 | 22 | 16 |
| Touch | Hand | Fingers_Hand \| Genitals | NA | NA | Touch.4 | 9 | 22 | 13 |
| Touch | Hand | Body | NA | NA | Touch.5 | 4 | 19 | 10 |
| Touch | Hand | Arm | NA | NA | Touch.6 | 6 | 22 | 9 |
| Touch | Fingers_Hand | Face_Mouth | NA | NA | Touch.7 | 20 | 20 | 3 |
| Touch | Hand | Head | NA | NA | Touch.8 | 1 | 26 | 4 |
| Touch | Hand | Face_Mouth | NA | NA | Touch.9 | 13 | 18 | 6 |
| * These Gesture Actions have been lumped with the non-repetitive version before performing the cluster analysis. Original Gesture Action definition includes repetition (i.e.,*Hitting Object*, *Hitting Other*, *Stomping Object*, *Stomping Other*). | | | | | | | | |

3. **Standard deviation of MAU and PAU durations**

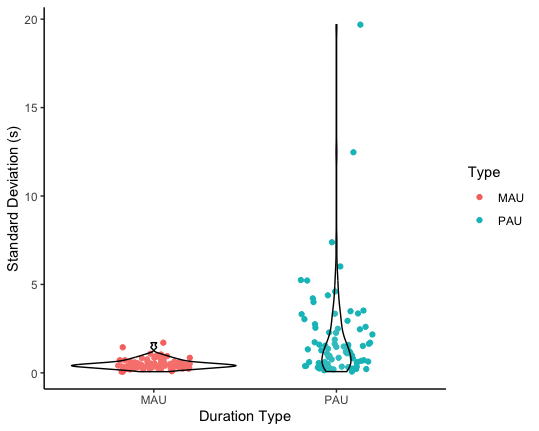

**Figure S3. Violin plots showing standard deviation distributions of MAU- and PAU-based durations for each Gesture Action across the three communities.** Points represent standard deviation values of duration of each Gesture Action type.

### 4. **Datasets**

Zipf’s law of brevity was tested on four datasets based on a) two levels of granularity and b) two units of duration. Specifically, the MAU-GA dataset contained the duration of the minimum action unit (MAU) of each token and the relative frequency of Gesture Actions (*rFq_action*) (level 1 granularity); the MAU-Morph dataset contained the duration of the MAU of each token and the relative frequency of gesture Morph *(rFq_morph)* (level 2 granularity); the PAU-GA dataset contained the duration of Performed Action Unit (PAU) of each token and the relative frequency of Gesture Action (*rFq_action*); PAU-Morph contained the duration of the PAU of each token and the relative frequency of gesture Morphs (*rFq_morph*) (Table S3). Because determining precise duration for each component was not always possible and because not all Gesture Actions were assigned a Morph, sample sizes per each subset vary (Table S3).

| **Table S3. Description of datasets employed in Zipf’s law of brevity analysis based on level of granularity and unit of duration.** | | |
| --- | --- | --- |
| **Granularity** | **Unit of duration** | |
|  | **MAU duration** | **PAU duration** |
| **Gesture Action** | MAU-GA dataset  (N=7267 tokens) | PAU-GA dataset  (N=7047 tokens) |
| **Morphs** | MAU-Morphs dataset  (N=7077 tokens) | PAU-Morph dataset  (N=6854 tokens) |

During data collection, three females emigrated from Sonso to Waibira and were present in both datasets.

5. **Model results**
   a. **Zipf-MAU-GA model**

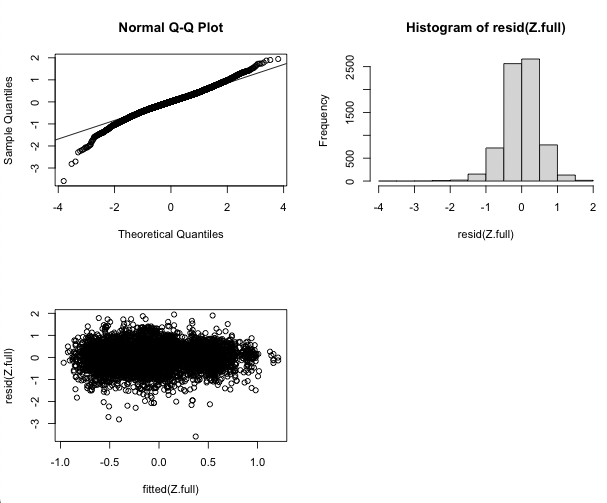
This model tested for the variation between communities in the relationship between Gesture Action relative frequency *rFq_action* and MAU duration. The VIFs for the Zipf-MAU-GA model having the log of the MAU duration as response variable and relative frequency of Gesture Action in interaction with community as fixed effects, with signaller, Gesture Action and goal as random factors were less than 1.08, indicating no issue of collinearity [7]. Model assessment showed no significant issue with heteroscedasticity or uneven residual distribution (Figure S4).

**Figure S4.** **QQ plot, histogram of residual distribution, and heterogeneity of variance graph for the Zipf-MAU-GA model, which assessed the effect of Gesture Action relative frequency *rFq_action* on the log of the MAU duration of gesture tokens**. In addition, the Zipf-MAU-GA model had signaller, Gesture Action, and goal as random factors.

| **Table S4. Summary of the results for the Zipf-MAU-GA model.** Reported are fixed factor estimates with standard errors, 95% confidence intervals, and p-values together with random effect values. Relevelled predictors indicate results from the same model but with changing reference level to allow for group-to-group comparisons. Significant results are highlighted in bold. | | | | |
| --- | --- | --- | --- | --- |
|  | **Log(MAU duration)** | | | |
| *Predictors* | *Estimates* | *Std. Error* | *95% CI* | *p-value* |
| (Intercept) | 1.21 | 0.07 | 1.07 – 1.34 | 0.001 |
| Gesture Action relative frequency *- rFq_action* | -1.22 | 0.70 | -2.60 – 0.16 | 0.083 |
| Community [Sonso] | *Reference* |  |  |  |
| Community [Waibira] | -0.01 | 0.04 | -0.10 – 0.07 | 0.729 |
| Community [Bossou] | -0.08 | 0.04 | -0.17 – 0.00 | 0.051 |
| **Community [Waibira - Bossou] - *Relevelled*** | **0.10** | **0.04** | **0.02 – 0.18** | **0.011** |
| *rFq_action* × Community [Sonso] | *Reference* |  |  |  |
| ***rFq_action*** **× Community [Waibira]** | **1.96** | **0.68** | **0.63 – 3.28** | **0.004** |
| *rFq_action* × Community [Bossou] | -0.20 | 0.93 | -2.03 – 1.63 | 0.832 |
| ***rFq_action*** **× Community [Waibira - Bossou] - *Relevelled*** | **1.61** | **0.71** | **0.22 – 3.00** | **0.023** |
| **Random Effects** | | | | |
| σ^2^ | 0.35 | | |  |
| Signaller (N=175) | 0.00 | | |  |
| Gesture Actions (N=87) | 0.25 | | |  |
| Goal (N=25) | 0.02 | | |  |
| ICC | 0.44 | | |  |
| Observations | 7267 | | |  |
| Marginal R^2^ / Conditional R^2^ | 0.008 / 0.441 | | |  |

The effect of *rFq_action* on the log MAU duration of Gesture Actions was significantly different between Sonso and Waibira (Estimate: 1.96 ± 0.68, 95% CI [0.63; 3.28]) and between Bossou and Waibira (Estimate: 1.61 ± 0.71, 95% CI [0.22; 3.00]).

Specifically, the effect of *rFq_action* on log MAU duration was steeper with increasing *rFq_action* in Waibira when compared to the other two communities (Table S4).

When assessing Zipf’s law of brevity within community only Bossou showed a significant negative effect of relative Gesture Action frequency *rFq_action* on gesture MAU duration (Tables S5, S6, S7).

| Table S5. Results for the Zipf-MAU-GA model computed on the Sonso subset which explored the effect of Gesture Action relative frequency *rFq_action* on the log of the MAU duration of gesture tokens. Reported are parameter estimates, standard error, 95% confidence intervals, and p-values together with random effect values. Significant results are highlighted in bold. | | | | |
| --- | --- | --- | --- | --- |
| **Sonso - Gesture Actions** | **Log(MAU duration)** | | | |
| *Predictors* | *Estimates* | *Std. Error* | *95% CI* | *p-value* |
| (Intercept) | 0.00 | 0.07 | -0.14 – 0.15 | 0.981 |
| Gesture Action relative frequency - *rFq_action* | -0.26 | 2.49 | -5.13 – 4.62 | 0.918 |
| **Random Effects** | | | |  |
| σ^2^ | 0.22 | | |  |
| Signaller (N=72) | 0.00 | | |  |
| Gesture Action (N=70) | 0.17 | | |  |
| Goal (N=24) | 0.02 | | |  |
| ICC | 0.46 | | |  |
| Observations | 4247 | | |  |
| Marginal R^2^ / Conditional R^2^ | 0.000 / 0.459 | | |  |

| Table S6. Results for the Zipf-MAU-GA model computed on the Bossou subset which explored the effect of Gesture Action relative frequency *rFq_action* on the log of the MAU duration of gesture tokens. Reported are parameter estimates, standard error, 95% confidence intervals, and p-values together with random effect values. Significant results are highlighted in bold. | | | | |
| --- | --- | --- | --- | --- |
| **Bossou – Gesture Actions** | **Log(MAU duration)** | | | |
| *Predictors* | *Estimates* | *Std. Error* | *95% CI* | *p-value* |
| (Intercept) | -0.05 | 0.09 | -0.21 – 0.12 | 0.586 |
| **Gesture Action relative frequency - *rFq_action*** | **-6.27** | **2.95** | **-12.06 – -0.48** | **0.034** |
| **Random Effects** | | | |  |
| σ^2^ | 0.23 | | |  |
| Gesture Action (N=72) | 0.16 | | |  |
| Signaller (N=30) | 0.00 | | |  |
| Goal (N=24) | 0.03 | | |  |
| ICC | 0.46 | | |  |
| Observations | 1784 | | |  |
| Marginal R^2^ / Conditional R^2^ | 0.047 / 0.482 | | |  |

| Table S7. Results for the Zipf-MAU-GA model computed on the Waibira subset which explored the effect of Gesture Action relative frequency *rFq_action* on the log of the MAU duration of gesture tokens. Reported are parameter estimates, standard error, 95% confidence intervals, and p-values together with random effect values. Significant results are highlighted in bold. | | | | |
| --- | --- | --- | --- | --- |
| **Waibira – Gesture Actions** | **Log(MAU duration)** | | |  |
| *Predictors* | *Estimates* | *Std. Error* | *95% CI* | *p-values* |
| (Intercept) | -0.02 | 0.08 | -0.18 – 0.14 | 0.810 |
| Gesture Action relative frequency -  *rFq_action* | 0.91 | 1.75 | -2.52 – 4.35 | 0.603 |
| **Random Effects** | | | |  |
| σ^2^ | 0.20 | | |  |
| Signaller (N=76) | 0.00 | | |  |
| Gesture Action (N=58) | 0.21 | | |  |
| Goal (N=24) | 0.02 | | |  |
| ICC | 0.53 | | |  |
| Observations | 1236 | | |  |
| Marginal R^2^ / Conditional R^2^ | 0.012 / 0.549 | | |  |

b. **Zipf-MAU-Morph model**

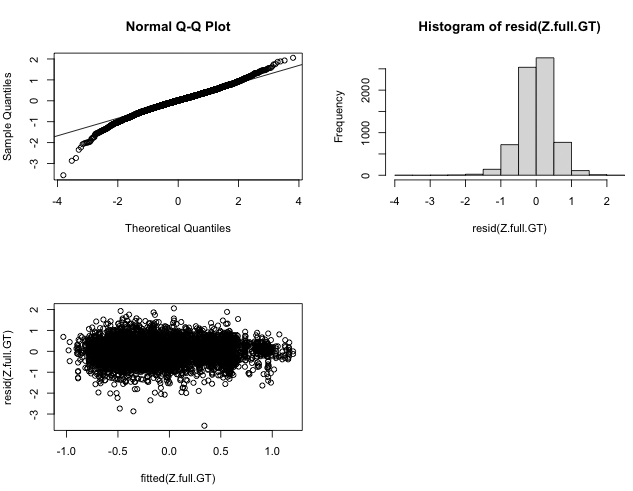
This model tested for the variation between communities in the relationship between Morph relative frequency *rFq_morph* and MAU duration. The Zipf-MAU-Morph model having the log of the MAU duration as response variable and relative frequency of Morph in interaction with community as fixed effects, with signaller, Gesture Action and goal as random factors had VIFs < 1.04, indicating no issue of collinearity [7]. Model assessment showed no significant issue with heteroscedasticity or uneven residual distribution (Figure S5).

**Figure S5. QQ plot, histogram of residual distribution, and heterogeneity of variance graph for the Zipf-MAU-Morph model, which assessed the effect of Morph relative frequency *rFq_morph* on the log of the MAU duration of gesture tokens**. In addition, the Zipf-MAU-Morph model had signaller, Morph, and goal as random factors.

The relationship between *rFq_morph* and log MAU was significantly different only between Sonso and Waibira, with the slope being steeper towards higher values for Waibira than for Sonso, (Table S8) while it did not differ between Sonso and Bossou (Table S8) or between Bossou and Waibira (Table S8; Figure S6). To assess the presence of a significant negative correlation in any of the three communities that would result in the expression of Zipf’s law of brevity, we ran the full model (excluding the community fixed factor) in subsets of the community-specific subsets of data.

Table S8. Summary of the results for the Zipf-MAU-Morph model. Reported are fixed factor estimates with standard errors, 95% confidence intervals, and p-values together with random effect values. Relevelled predictors indicate results from the same model but with changing reference level to allow for group-to-group comparisons. Significant results are highlighted in bold.

| Zipf-MAU-Morph | **Log(MAU duration)** | | | |
| --- | --- | --- | --- | --- |
| *Predictors* | *Estimates* | *Std. Error* | *95% CI* | *p-value* |
| (Intercept) | -0.05 | 0.04 | -0.13 –0.04 | 0.282 |
| Morph relative frequency - *rFq_morph* | 0.71 | 0.85 | -0.97 – 2.38 | 0.407 |
| Community [Sonso] | *Reference* |  |  |  |
| Community [Waibira] | -0.06 | 0.03 | -0.13 – -0.00 | 0.049 |
| Community [Bossou] | -0.19 | 0.03 | -0.25 – -0.13 | <0.001 |
| Community [Waibira – Bossou] – *Relevelled* | 0.13 | 0.13 | 0.05 – 0.20 | 0.001 |
| *rFq_morph* × Community [Sonso] | *Reference* |  |  |  |
| ***rFq_morph* × Community [Waibira]** | **2.66** | **2.66** | **0.84 – 4.47** | **0.004** |
| *rFq_morph* × Community [Bossou] | -1.90 | 1.17 | -0.38 – 4.18 | 0.103 |
| *rFq_morph* × Community [Waibira – Bossou] – *Relevelled* | 0.76 | 1.22 | -1.63 – 3.14 | 0.536 |
| **Random Effects** | | | | |
| σ^2^ | 0.24 | | | |
| Signaller (N=174) | 0.00 | | | |
| Morph (N=140) | 0.14 | | | |
| Goal (N=25) | 0.01 | | |  |
| ICC | 0.40 | | |  |
| Observations | 7077 | | |  |
| Marginal R^2^ / Conditional R^2^ | 0.017 / 0.407 | | |  |

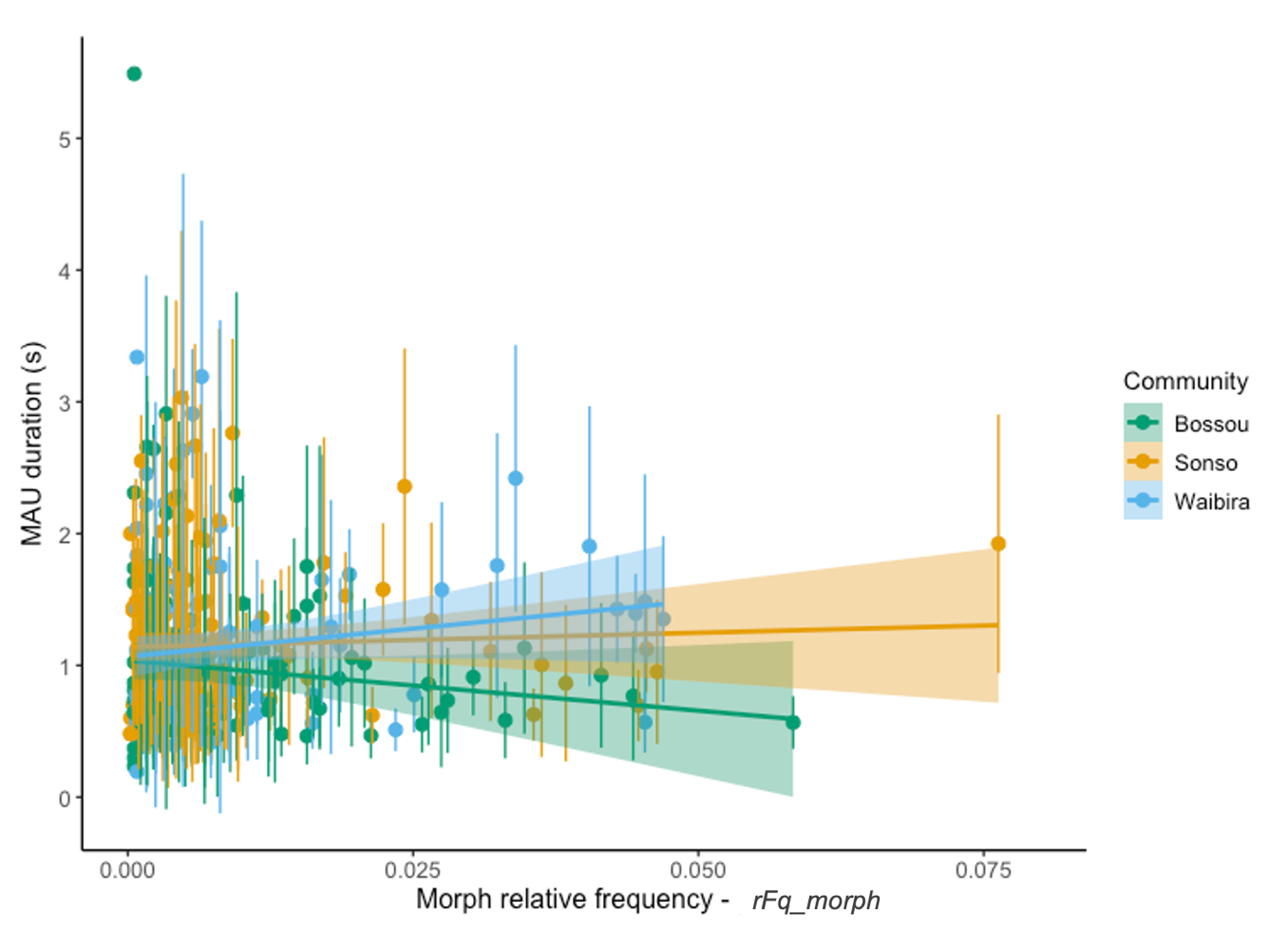
When assessing Zipf’s law of brevity within community no community had a significant negative effect of relative Morph frequency *rFq_morph* on gesture MAU duration (Tables S9, S10, S11).

**Figure S6. Scatterplot with regression lines representing the relationship between the MAU duration of Morph and their relative frequency (*rFq_morph*).** Points indicate Morphs, bars indicate standard error, colours indicate community, shaded area indicates 95% confidence intervals of regression line.

| Table S9. Results for the Zipf-MAU-Morph model computed on the Sonso subset which explored the effect of Morph relative frequency *rFq_morph* on the log of the MAU duration of gesture tokens. Reported are parameter estimates, standard error, 95% confidence intervals, and p-values together with random effect values. Significant results are highlighted in bold. | | | | |
| --- | --- | --- | --- | --- |
| **Sonso - Morph** | **Log(MAU duration)** | | | |
| *Predictors* | *Estimates* | *Std. Error* | *95% CI* | *p-value* |
| (Intercept) | -0.05 | 0.06 | -0.16 – 0.06 | 0.417 |
| Morph relative frequency – *rFq_morph* | 1.77 | 3.27 | -4.64 – 8.18 | 0.589 |
| **Random Effects** | | | |  |
| σ^2^ | 0.23 | | |  |
| Morph (N=127) | 0.16 | | |  |
| Signaller (N=72) | 0.00 | | |  |
| Goal (N=24) | 0.02 | | |  |
| ICC | 0.43 | | |  |
| Observations | 4187 | | |  |
| Marginal R^2^ / Conditional R^2^ | 0.003 / 0.436 | | |  |

| Table S10. Results for the Zipf-MAU-Morph model computed on the Waibira subset which explored the effect of Morph relative frequency *rFq_morph* on the log of the MAU duration of gesture tokens. Reported are parameter estimates, standard error, 95% confidence intervals, and p-values together with random effect values. Significant results are highlighted in bold. | | | | |
| --- | --- | --- | --- | --- |
| **Waibira - Morph** | **Log(MAU duration)** | | | |
| *Predictors* | *Estimates* | *Std. Error* | *95% CI* | *p-values* |
| (Intercept) | -0.11 | 0.07 | -0.25 – 0.02 | 0.099 |
| Morph relative frequency -  *rFq_morph* | 4.89 | 2.84 | -0.69 – 10.46 | 0.086 |
| **Random Effects** | | | |  |
| σ^2^ | 0.20 | | |  |
| Morph (N=102) | 0.17 | | |  |
| Signaller (N=75) | 0.01 | | |  |
| Goal (N=24) | 0.02 | | |  |
| ICC | 0.50 | | |  |
| Observations | 1180 | | |  |
| Marginal R^2^ / Conditional R^2^ | 0.046 / 0.525 | | |  |

| **Table S11. Results for the Zipf-MAU-Morph model computed on the Bossou subset which explored the effect of Morph relative frequency *rFq_morph* on the log of the MAU duration of gesture tokens**. Reported are parameter estimates, standard error, 95% confidence intervals, and p-values together with random effect values. Significant results are highlighted in bold. | | | | |
| --- | --- | --- | --- | --- |
| **Bossou - Morph** | **Log(MAU duration)** | | | |
| *Predictors* | *Estimates* | *Std. Error* | *95% CI* | *p-values* |
| **(Intercept)** | **-0.19** | **0.07** | **-0.32 – -0.06** | **0.005** |
| Morph relative frequency - *rFq_morph* | -4.35 | 3.86 | -11.92 – 3.21 | 0.259 |
| **Random Effects** | | | |  |
| σ^2^ | 0.26 | | |  |
| Morph (N=126) | 0.14 | | |  |
| Signaller (N=30) | 0.00 | | |  |
| Goal (N=24) | 0.03 | | |  |
| ICC | 0.40 | | |  |
| Observations | 1710 | | |  |
| Marginal R^2^ / Conditional R^2^ | 0.010 / 0.410 | | |  |

c. **Zipf-PAU-GA model**

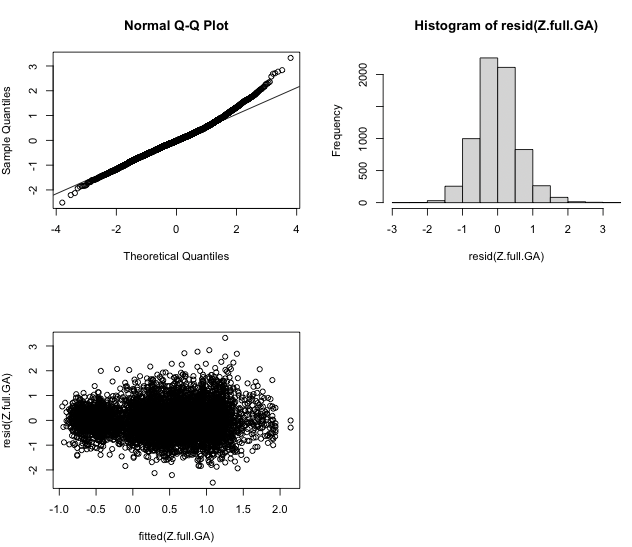
This model tested for the variation between communities in the relationship between Gesture Action relative frequency *rFq_action* and PAU duration. The VIFs for the Zipf-PAU-Morph model having the log of the performed action unit (PAU) duration as response variable and relative frequency of Gesture Actions in interaction with community as fixed effects, with signaller, Gesture Action and goal as random factors were < 1.08, indicating no issue of collinearity [7]. Model assessment showed no significant issue with heteroscedasticity or uneven residual distribution (Figure S7).

**Figure S7. QQ plot, histogram of residual distribution, and heterogeneity of variance graph for the Zipf-PAU-GA model, which assessed the effect of Gesture Action relative frequency *rFq_action* on the log of the Gesture duration of gesture tokens**. In addition, the Zipf-PAU-GA model had signaller, Gesture Action, and goal as random factors.

Because the interaction factor of relative Gesture Action frequency *rFq_action* and community did not improve model fit (assessed using the ‘drop1’ function; LRT: X^2^=4.83, p=0.09), we ran the model once more without the interaction to allow for the assessment of individual fixed effects. The lack of a significant interaction suggested an overarching pattern across communities and indicated no specific community variation. Overall, there was no significant effect of *rFq_action* on the log of the PAU duration, suggesting a lack of Zipf’s law of brevity across all communities in the dataset. The only significant difference detected was the in the PAU durations across communities, with Bossou and Waibira performing shorter Gesture Actions across the repertoire, as compared to Sonso (Table S12, Figure S8, Figure S9).

| **Table S12. Summary of the results for the Zipf-PAU-GA model.** Reported are fixed factor estimates with standard errors, 95% confidence intervals, and p-values together with random effect values. Relevelled predictors indicate results from the same model but with changing reference level to allow for group-to-group comparisons. Significant results are highlighted in bold. | | | | |
| --- | --- | --- | --- | --- |
|  | **Log(PAU duration)** | | | |
| *Predictors* | *Estimates* | *Std. Error* | *95% CI* | *p-value* |
| (Intercept) | 0.36 | 0.08 | 0.20 – 0.52 | <0.001 |
| Gesture Action relative frequency - *rFq_action* | 0.09 | 0.31 | -0.51 – 0.69 | 0.759 |
| Community [Sonso] | *Reference* |  |  |  |
| **Community [Waibira]** | **-0.14** | **0.03** | **-0.19 – -0.08** | **<0.001** |
| **Community [Bossou]** | **-0.19** | **0.03** | **-0.25 – -0.14** | **<0.001** |
| Community [Waibira - Bossou] - *Relevelled* | 0.06 | 0.03 | -0.01 – 0.12 | 0.088 |
| **Random Effects** | | | |  |
| σ^2^ | 0.36 | | |  |
| Signaller (N=174) | 0.00 | | |  |
| Gesture type (N=87) | 0.41 | | |  |
| Goal (N=25) | 0.02 | | |  |
| ICC | 0.55 | | |  |
| Observations | 7047 | | |  |
| Marginal R^2^ / Conditional R^2^ | 0.009 / 0.555 | | |  |

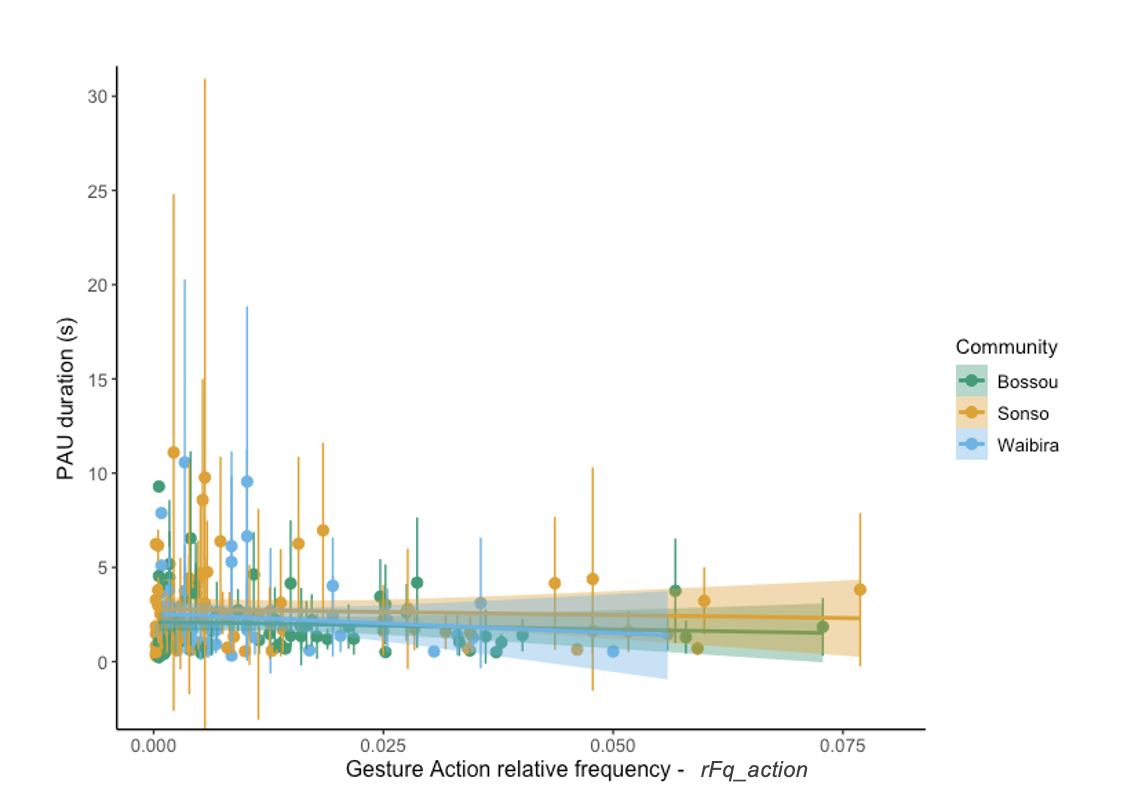

**Figure S8. Scatterplot with regression lines representing the relationship between the PAU duration of Gesture Actions and their relative frequency (*rFq_action*).** Points indicate Gesture Actions. Bars indicate standard error, colours indicate community, shaded area indicates 95% confidence intervals of regression line.

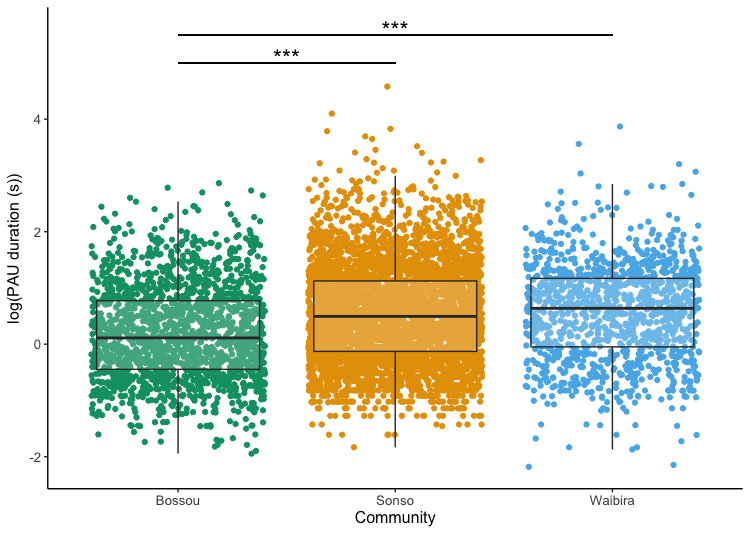
**Figure S9. Boxplot with jittered points representing the difference in PAU duration across communities for the Gesture Action subset**. Points indicate gesture tokens, colours indicate community. Statistically significant differences in gesture PAU durations between community levels are marked with asterisks (***). Negative log values indicate durations of length between 0s and 1s.

d. **Zipf-PAU-Morph model**

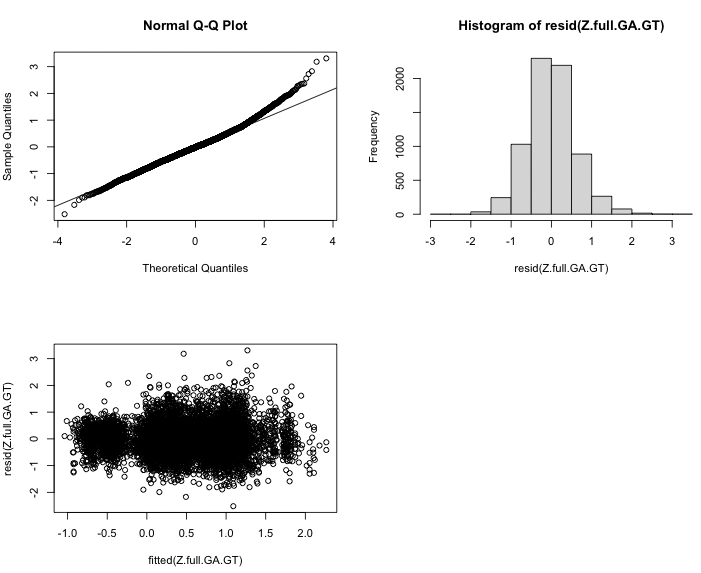
This model tested for the variation between communities in the relationship between Morph relative frequency *rFq_morph* and MAU duration. The VIFs for the Zipf-PAU-Morph model having the log of the performed action unit (PAU) duration as response variable and relative frequency of Morph in interaction with community as fixed effects, with signaller, Gesture Action and goal as random factors were < 1.04, indicating no issue of collinearity [7]. Model assessment showed no significant issue with heteroscedasticity or uneven residual distribution (Figure S10).

**Figure S10. QQ plot, histogram of residual distribution, and heterogeneity of variance graph for the Zipf-PAU-Morph model, which assessed the effect of Morph relative frequency *rFq_morph* on the log of the Gesture duration of gesture tokens**. In addition, the Zipf-PAU-Morph model had signaller, Morph, and goal as random factors.

Because the interaction factor of relative Morph relative frequency *rFq_morph* and community did not improve model fit (assessed using the ‘drop1’ function; X^2^=53.84, p=0.14), we ran the model once more without the interaction to allow for the assessment of individual fixed effects. The lack of a significant interaction suggested an overarching pattern across communities and indicated no specific community variation. The relative Morph frequency did not have any effect on the log PAU duration (Table S13), suggesting an absence of Zipf's law of brevity in this unit of analysis. Similarly to the analysis on Gesture Actions, Sonso showed longer PAU durations overall, as compared to those in both Bossou and Waibira (Table S13, Figure S11).

| **Table S13. Summary of the results for the Zipf-PAU-Morph model.** Reported are fixed factor estimates with standard errors, 95% confidence intervals, and p-values together with random effect values. Relevelled predictors indicate results from the same model but with changing reference level to allow for group-to-group comparisons. Significant results are highlighted in bold. | | | | |
| --- | --- | --- | --- | --- |
| Zipf-PAU-Morph | **Log(PAU duration)** | | | |
| *Predictors* | *Estimates* | *Std. Error* | *95% CI* | *p-values* |
| (Intercept) | 0.57 | 0.06 | 0.45 – 0.69 | <0.001 |
| Morph relative frequency – *rFq_morph* | 1.18 | 0.73 | -0.26 – 2.61 | 0.108 |
| Community [Sonso] | *Reference* |  |  |  |
| **Community [Waibira]** | **-0.16** | **0.03** | **-0.22 – -0.10** | **<0.001** |
| **Community [Bossou]** | **-0.21** | **0.03** | **-0.27 – -0.15** | **<0.001** |
| Community [Waibira - Bossou] - *Relevelled* | 0.05 | 0.04 | -0.02 – 0.12 | 0.162 |
| **Random Effects** | | | |  |
| σ^2^ | 0.37 | | |  |
| Signaller (N=173) | 0.01 | | |  |
| Morph (N=140) | 0.34 | | |  |
| Goal (N=25) | 0.03 | | |  |
| ICC | 0.50 | | |  |
| Observations | 6854 | | |  |
| Marginal R^2^ / Conditional R^2^ | 0.013 / 0.507 | | |  |

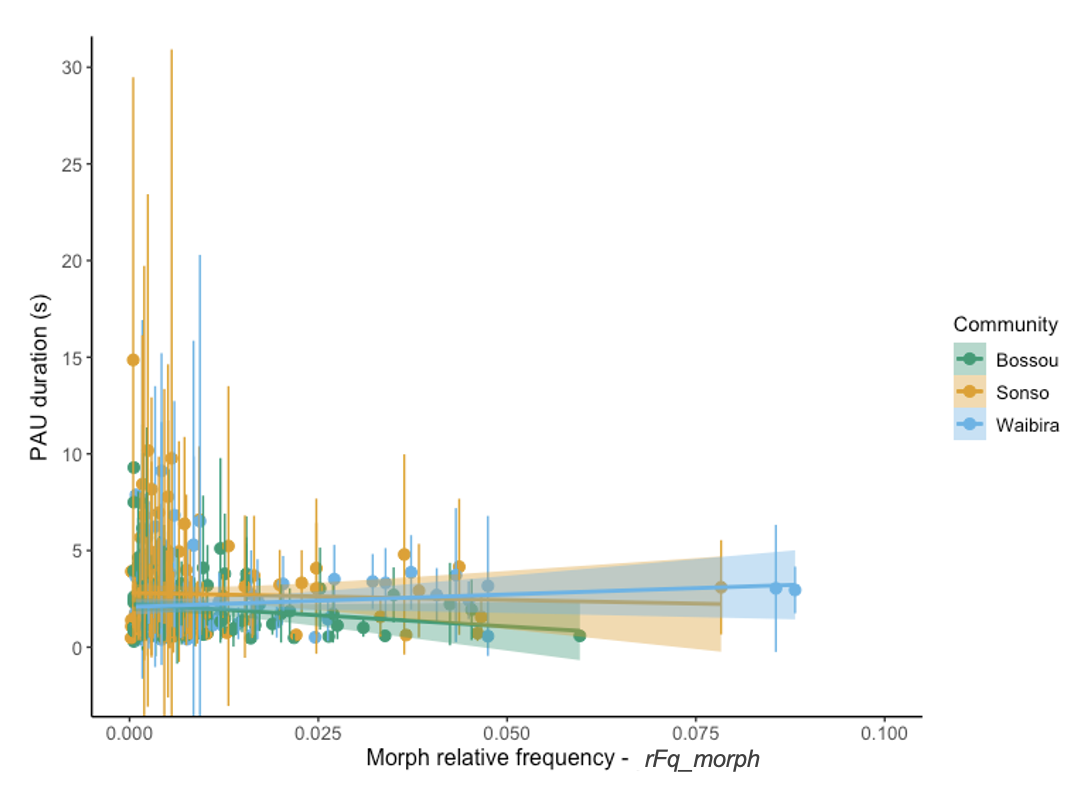
**Figure S11. Scatterplot with regression lines representing the relationship between the PAU duration of Morphs and their relative frequency (*rFq_morph*).** Points indicate Morphs, bars indicate standard error, colours indicate community, shaded area indicates 95% confidence intervals of regression line.

6. **Video data collection**

For Sonso, videos were collected between November 2007 and August 2012, between February 2017 and March 2022 for Waibira, and between January 1992 and December 2017 for Bossou. For the East African communities, video observations were collected between 7.30 and 16.30 with recording of gestures following ad libitum sampling [8]. Any social interaction that could lead to gestural communication was recorded (i.e., any situation involving two or more chimpanzees in proximity and not involved in solitary activities). If multiple opportunities for gestural data collection happened simultaneously, preference was given to those individuals from whom fewer data had been collected.

For Sonso chimpanzees, from October 2007 to August 2009, a Sony Handycam (DCR-HC-55) was used. Videos were recorded using MiniDV tape. Due to the challenges of filming wild chimpanzees in a visually dense rainforest, there were instances when the beginning of gestural sequences was not captured on video. In such cases, the missing parts were dictated at the end of the video, and these sequences were not included in the analysis.

From 2009 onwards in Sonso, and for all data collection in Waibira, video data were collected using Panasonic camcorders (V770, HC-VXF1), which have a 3-second pre-record feature that improves the ability to capture the start of behaviour. The same procedure was followed, and any sequences where the initiation of gesturing was not clearly captured continued to be excluded. Data collection for the Budongo chimpanzees was approved by the University of St Andrews Committee for Animal Welfare and Ethics.

For Bossou, videos were collected between 7.00 and 18.00 between the months of November and March during dry seasons in this region. Video recordings of the Bossou community were collected at two ‘outdoor laboratories’ located in the chimpanzees’ home range, established in 1989 and 2009 to study their tool use: called as Bureau and Salon [9]. Chimpanzees were filmed using two or three video cameras (mainly Sony Handycams such as DCR-PC 110 or DCR-VX 1000) originally recorded on VHS, Betamax, Hi8, MiniDV and digitized during the Bossou Video Archive project funded by JSPS Core-to-Core program (CCSN) and Cooperative Research program of Primate Research Institute, Kyoto University. Both Bureau and Salon are flat and clear of vegetation, allowing researchers to record chimpanzee behaviour clearly from a viewing station behind a vegetation screen with holes for filming, placed at around 15-20m from the centre of the labs. In addition, the lack of vegetation allows for accurate tracking of all individuals joining or leaving the party as well as recording of most communicative signals produced.

7. **Inter-observer reliability**

Reliability of coding accuracy was assessed by conducting ICC, Cohen’s Kappa, or percentage agreement on selected variables, on a subset of the data, representing around 5% of the total gestures coded. For the East African chimpanzees, IOR was carried as part of the establishment of an East African chimpanzee gesture database, which comprises gestures from five different communities (Sonso and Waibira in Budongo, Uganda; Issa in Issa Valley, Tanzania; Kanyawara in Kibale Forest, Uganda; and Kalinzu in Kalinzu Foreset, Uganda). First, a list of the Gesture Actions present at least 10 times across these five communities was established. Then, per community, a list of the total frequency of these Gesture Actions was created. A random sampling function was written to select 5% of the frequency of each Gesture Action within each community and create the sample for IOR testing. We capped the frequency of specific Gesture Actions within the 5% sample at seven examples, and we included at least one example of each Gesture Action. Within the larger project of establishing an East African gesture database, a second coder Gal Badihi (GB) re-coded gestures which were originally collected by ASa, and vice versa. A total of 320 gestures were re-coded belonging to 289 communications (AS coded 181 tokens from 168 communications; GB coded 139 tokens from 121 communications). Agreement assessment on Gesture Action, Gesture Action modifiers, durations, and communication tiers was carried out independently.

Overall agreement between the two coders for the East African dataset (ASa, GB) was excellent for both MAU duration (ICC= 1) and PAU duration (ICC=1). There was substantial agreement on the identification of Gesture Actions (Cohen’s Kappa=0.798), and agreement on the modifiers employed in the clustering analysis for Morph creation was near perfect (Repetition (Y/N) Cohen’s Kappa=0.952; Body part, Cohen’s Kappa=0.856). Inter observer agreement regarding the intentionality criteria markers was also good. The original coding categories in the data were lumped and, given a strong bias towards coding intentional gestures, we used percentage agreement (Gaze before: 91% agreement; Gaze during: 92.4% agreement, Directedness towards recipient: 94.07% agreement). Agreement on communicative goal was substantial (Cohen’s Kappa=0.786).

A similar procedure was employed for the Bossou chimpanzees, where a 5% subset of gesture examples was created for each dataset coded by ASa and a second coder Daniela Rodrigues (DR), making a total of 149 tokens. ASa coded 59 tokens from 53 communications, originally coded by DR; while DR coded 90 tokens from 84 communications, originally coded by AS.

Overall agreement between the two coders for the West African dataset (ASa, DR) was again excellent for both MAU duration (ICC= 1) and PAU duration (ICC=1). There was also almost perfect agreement on the identification of Gesture Actions (Cohen’s Kappa=0.801), and the other modifiers employed in the clustering analysis for Morph creation matched substantially (Repetition (Y/N) Cohen’s Kappa=0.699; Body part, Cohen’s Kappa = 0.793). Inter observer agreement regarding the intentionality criteria markers was also good (Gaze before = 87.3% agreement; Gaze during = 94.3% agreement) and substantial (Directedness towards recipient, Cohen’s Kappa = 0.799). Agreement on communicative goal was almost perfect (Cohen’s Kappa = 0.874).
